# Genomic analysis of laboratory-evolved, heat-adapted *Escherichia coli* strains

**DOI:** 10.1101/2024.10.01.616104

**Authors:** Bailey E. McGuire, Francis E. Nano

**Affiliations:** Department of Biochemistry and Microbiology, University of Victoria, Victoria, B.C. Canada

**Keywords:** Adaptive laboratory evolution (ALE), Heat shock protein, Thermotolerance, Adaptation to heat, Evolutionary genomics

## Abstract

Adaptive laboratory evolution to high incubation temperatures represents a complex evolutionary problem, and each study to date performed in *Escherichia coli* has resulted in a different set of mutations. We performed adaptive laboratory evolution of *E. coli* to heat by passaging a culture at elevated temperatures for 150 days. Throughout the adaptive evolution we expressed a set of genes that induce hyper-mutagenesis. These growth conditions yielded a strain with a maximum growth temperature approximately 2 °C above that of the parental strain. We preserved evolved isolates weekly and obtained and analyzed whole-genome sequencing data for three isolates from different time points. We identified hundreds of mutations, including mutations in components of the RNA polymerase (RpoB, RpoC and RpoD), Rho, and the heat shock proteins GroES, GroEL, DnaK, ClpB, IbpA and HslU. We compared the proteomes of the starting strain and final strain grown at 37 °C and 42.5 °C and identified changes in abundance between samples for GroESL, HslVU, DnaK, ClpB and other important proteins. This study details a distinct evolutionary route towards enhanced thermotolerance, contributes to our understanding of adaptation to heat in *Escherichia coli* and may provide insights into heat adaptation in other organisms.

## INTRODUCTION

Temperature affects all molecules in a cell, and as such, changes in growth temperature parameters like minimums, optimums and maximums represent complex evolutionary problems. Cells also have adapted to survive temporary exposures to extremes of heat and cold. A better understanding of how organisms evolve to changes in environmental temperatures will advance our knowledge about how organisms are evolving to the challenges of climate change and may allow us to engineer important organisms for better survival in warming environments. One such application currently being pursued is in evolving thermotolerance in corals and their associated microbes in an effort to prevent the loss of coral reefs^1^.

In the laboratory, researchers can simulate many aspects of evolution, and when they do so under specific conditions this is often termed adaptive laboratory evolution (ALE). ALE can be thought of as genome-wide directed evolution, where the diversification, screening and selection steps are carried out simultaneously by selecting for strains that have superior growth in a defined set of conditions. Rather than relying on the spontaneous mutation rate of the organism, diversification in ALE studies is sometimes accelerated through hyper-mutagenesis. To this end, some studies have employed stationary phase mutagenesis or overexpressed a protein which raises the mutation rate of the cell^2,3^. Depending on the hyper-mutagenesis method, certain kinds of genetic changes will likely occur more or less frequently than during natural evolution. Microorganisms are amenable to ALE due to their short doubling times. In a typical ALE experiment cells are serially cultured in the presence of a stressor, and the severity of that stress is increased over time. In previous *E. coli* high-temperature ALE studies, researchers periodically increased the incubation temperature while subculturing the cells. Upon whole-genome sequencing (WGS) analysis, they often found changes in master regulator, stress response and other classes of genes in the evolved isolates^2–7^. In related studies performed at constant temperatures above the *E. coli* optimum, changes in master regulator genes were shown to cause changes in the expression of hundreds of genes^8–11^. Changes in stress response genes could cause changes in gene expression as well, perhaps differently under stressful versus not stressful conditions.

In this study, we used an inducible mutagenesis plasmid, MP6^12^, to enhance the ALE of *E. coli* to high incubation temperatures. We evolved the final strain for 150 days, and we analyzed the final strain, the starting strain and two intermediate isolates by WGS. We observed increases in the maximum growth temperature and ethanol resistance of the lineage during the adaptation to growth at higher temperatures. The mutations that accumulated in the heat-resistant strains represent an evolutionary pathway that has not been previously observed. Namely, our strain contains unique amino acid substitutions in RpoB, RpoC and Rho, and unlike other strains from similar studies, there are many amino acid substitutions in DNA topoisomerase, DNA gyrase and various heat shock proteins. Finally, we analyzed the proteomes of the starting DH10B strain and the final heat-evolved isolate HE_150 each grown at 37 °C and 42.5 °C, in duplicate.

## RESULTS

### Generating heat-evolved strains with ALE to heat and MP6 mutagenesis

The widely-used *E. coli* laboratory strain DH10B harboring plasmid MP6^12^ was subjected to ALE through growth at increasing temperature while inducing the mutagenic DNA replication, repair and base selection proteins encoded by MP6. The *E. coli* strain was serially cultured, and the incubation temperature was increased periodically, starting at 43.5 °C and ending at 46.9 °C (**Fig. 1**). When induced, the MP6 plasmid will affect mutagenesis of itself as well as the chromosome; thus, at different time points we cured MP6 and re-introduced parental MP6 from a master stock (**Fig. 1**). Cells were designated HE_X, indicating <u>H</u>eat-<u>E</u>volved cells that experienced <u>X</u> 24-hour growth cycles in the presence of 0.1 mM arabinose inducer, which induces gene expression from plasmid MP6. Select HE isolates are available through Addgene: HE_50 (Addgene ID 201049), HE_100 (Addgene ID 201050) and HE_150 (Addgene ID 201051).

**Figure 1.**
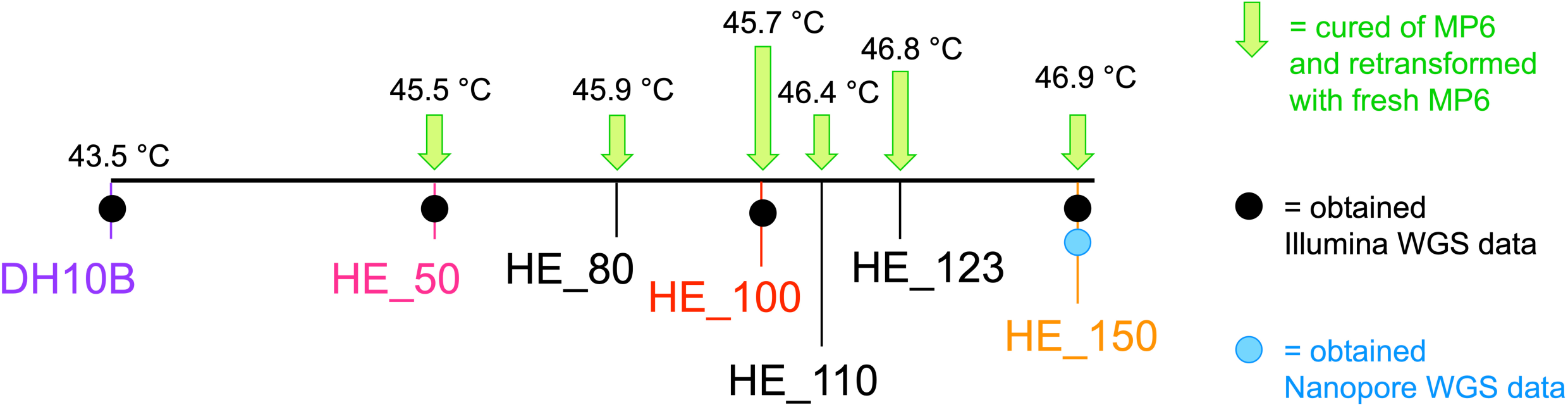
Heat evolution experimental details. Schematic showing events over the heat evolution experiment, including average incubator temperatures, timepoints where isolates were cured of their MP6 plasmids and retransformed with fresh MP6, names of select isolates, and isolates which were analyzed via WGS.

### ALE to heat shifted the growth parameters of HE strains

Growth experiments of the parental DH10B strain and HE strains revealed clear differences. Colony formation on agar medium provided a clear-cut growth/no growth distinction, and we used colony formation as evidence of viability at a given temperature. On agar plates, parental DH10B formed some isolated colonies at 45 °C, HE_50 and HE_100 formed isolated colonies at 45 and 46.8 °C, and HE_150 formed isolated colonies at 45, 46.8 and 47 °C (**Fig. 2**). Thus, the final HE isolate HE_150 has a maximum plate growth temperature approximately 2 °C above that of the starting DH10B strain.

**Figure 2.**
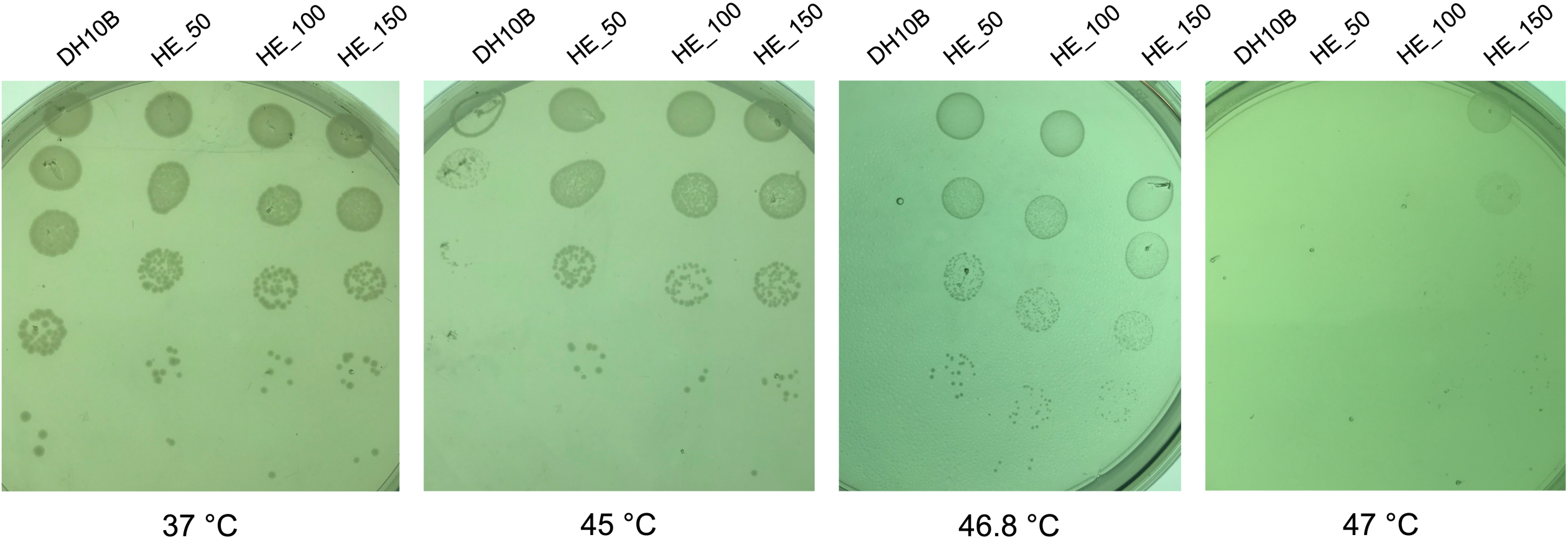
Plate growth of parental DH10B and HE cells at high temperatures. Cell suspensions at 1 OD_600_ unit were serially diluted and 10 µL of dilutions from 10^-2^ to 10^-6^ were spotted onto LB agar. The plates were incubated at the indicated temperatures for 24 hours.

We measured the growth of DH10B, HE_50, HE_100 and HE_150 (**Fig. 3**) in broth cultures at 37.3 °C, 41.3 °C and 45.3 °C. At 45.3 °C, DH10B cultures reached stationary phase towards the end of the 320-minute time period (**Fig. 3A** and **3E**), and the cultures did not significantly increase in optical density upon further incubation (**Supplementary Fig. S1**). Previous studies have observed an increased resistance to heat when strains were grown on agar media rather than in liquid media^2,4^, increases in optical densities from inclusion body formation^13^ and the ability to calculate doubling times for cells grown at lethal temperatures^4^, complicating growth assessment at high temperatures. Thus, increasing optical density values for parental DH10B at 45.3 °C don’t necessarily indicate growth. In contrast, HE growth at 45.3 °C showed less leveling off (**Fig. 3B-E**), and the cultures significantly increased in optical density when incubated past the 320-minute time point (**Supplementary Fig. S1**). At the 23-hour time point, the 45.3 °C optical densities of the HE cultures were significantly higher than those of the DH10B cultures (**Supplementary Fig. S2A**). We also grew the four strains at 46.3 °C for 23 hours and observed a more dramatic difference in optical densities between parental DH10B and HE cells (**Supplementary Fig. S2B**).

**Figure 3.**
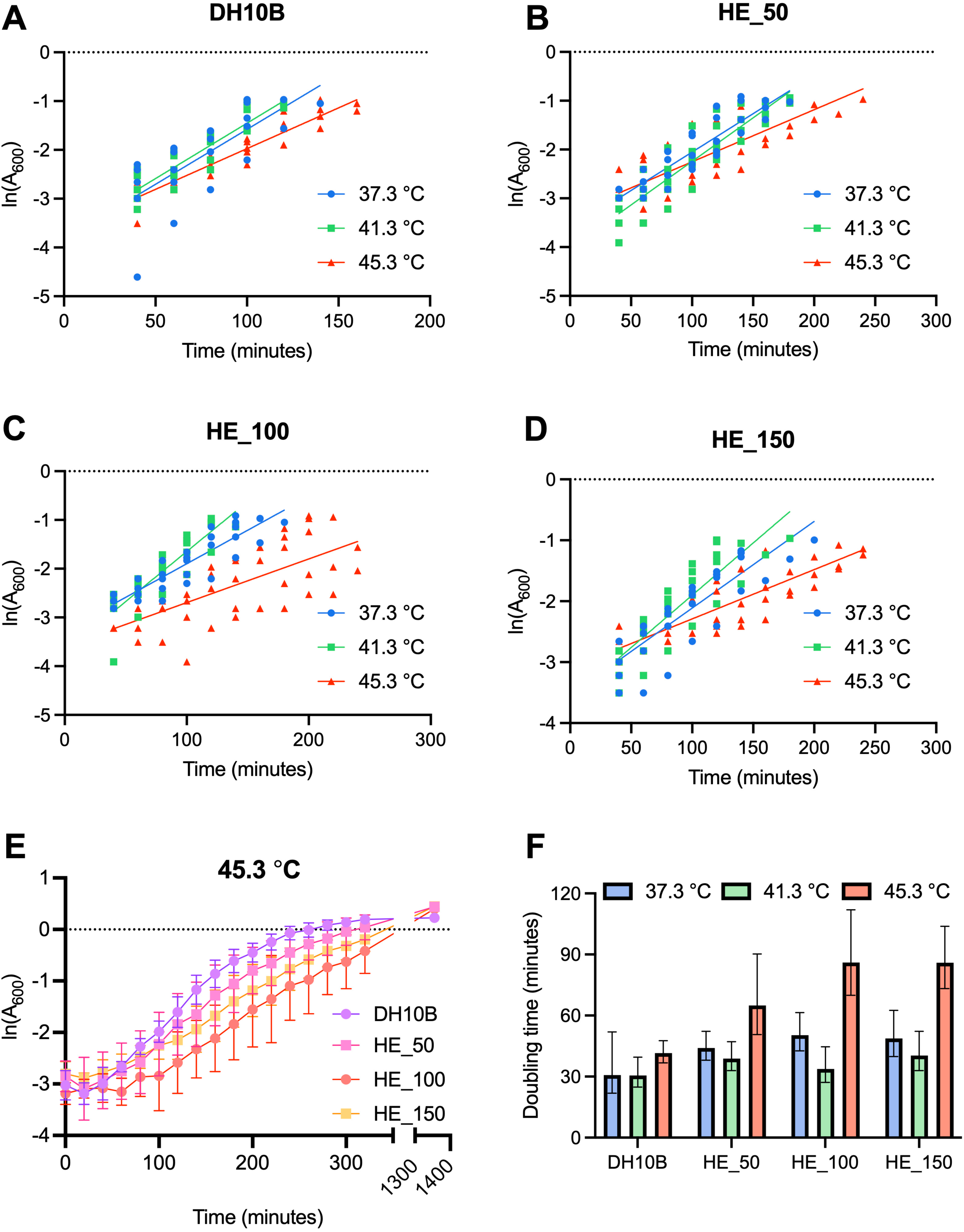
Liquid culture growth characteristics of parental DH10B and HE cells at high temperatures. **A-D.** Parental DH10B (**A**), HE_50 (**B**), HE_100 (**C**) and HE_150 (**D**) semi-log growth curves of the exponential growth phase at 37.3 °C, 41.3 °C and 45.3 °C. The absorbance at 600 nm of each strain at each temperature was measured every 20 minutes in quintuplicate. The ln(A_600_) values corresponding to A_600_ values ≤0.4 were used and time points 0 and 20 minutes were removed from the analysis to exclude the lag phase. **E.** Semi-log growth curves of of parental DH10B and HE cultures at 45.3 °C. The absorbance was measured at 20-minute intervals from time 0 to 320 minutes (5 hours and 20 minutes) and again at 1355 minutes (22 hours and 35 minutes). Error bars represent the standard deviation from the mean. **F.** Doubling times of parental DH10B and HE cells at 37.3, 41.3 and 45.3 °C, calculated from the semi-log graphs in **A-D**. Error bars represent the 95 % confidence interval.

Comparison of the growth curves suggest that ALE to heat shifted the optimum growth temperature (**Fig. 3A-E**). The calculated doubling times for each strain at each temperature demonstrate differences between parental DH10B and HE cells (**Fig. 3F**), including the observation that doubling times for parental DH10B at 37.3 °C and 41.3 °C were more similar whereas doubling times for HE strains trend towards being shorter at 41.3 °C than 37.3 °C. The doubling times of HE cells tended to be longer than those of parental DH10B at most temperatures (**Fig. 3F**), perhaps due to fixation of deleterious mutations alongside beneficial and neutral ones.

### ALE to heat induced resistance to ethanol

Throughout their heat evolution, HE isolates appeared to acquire phenotypic changes besides resistance to increased temperatures. One phenotype that commonly accompanies an increased resistance to elevated temperatures is increased resistance to organic solvents such as ethanol. Indeed, when we tested the resistance of HE_50, HE_100 and HE_150, we found that all three strains exhibited substantial growth in the presence of 4 % ethanol, whereas DH10B was unable to grow in these conditions (**Supplementary Fig. S3**).

### WGS of heat evolved cells

Chromosomal DNA from strains DH10B, HE_50, HE_100 and HE_150 were subjected to Illumina WGS; HE_150 DNA was also analyzed by Oxford Nanopore WGS. Two to 2.8 million trimmed paired-end Illumina reads were mapped to the appropriate reference genomes, resulting in mean coverages of 58-76 per bp. Oxford Nanopore sequencing of HE_150 generated 82,131 reads, with a mean length of 5 kb and a maximum length of 65 kb, representing a mean coverage of 89 per base. Additional WGS statistics can be found in **Supplementary Tables S1** and **S2**. We built the parental DH10B and HE isolate genomes through a combination of mapping the reads to the DH10B reference genome (CP000948.1), SPAdes *de novo* assembly of the HE_150 genome and manual curation. We submitted the raw WGS reads to the SRA with the accession numbers SRR21983746-SRR21983750. We also submitted the genome sequences of our DH10B isolate (CP110018), HE_50 (CP110017), HE_100 (CP110016) and HE_150 (CP110015) to Genbank.

Both the reference DH10B genome and our isolate of DH10B have a ∼227 kb region which is a tandem duplication of 113 kb of the chromosome. When we mapped Illumina reads to an HE draft genome with just half of the repeat region, we identified mutations in the region. The number of mutations in this region corresponded well to the number expected in a region of that length for all HE strains (**Supplementary Table S3**). We did not include these mutations in the final genomes since we could not determine in which half of the repeat region each mutation occurred. Thus, we include these variations in **Supplementary File S2**.

### ALE to heat produced hundreds of genetic changes

We discovered hundreds of genomic changes that arose during the heat evolution process and a few mutations which were lost between early and late HE strains (**Supplementary Fig. S4** and **Supplementary File S2**). Throughout their heat evolution, the HE strains acquired 7-11 changes on average in every 24-hour period (**Supplementary Table S4**). For comparison, if *E. coli* had a spontaneous mutation rate on the higher end of normal of one change in every billion base pairs per generation and a doubling time of one hour, we would expect ∼0.1 changes in a 24-hour period. Single nucleotide polymorphisms (SNPs) accounted for 87-88 % of the changes in HE isolates, meaning on average, 6-10 SNPs arose every 24 hours. The remaining changes included small indels and the acquisition of insertion sequence elements, including the insertion of an IS186B element in *sibB* in our DH10B strain and all the HE strains and an IS1A insertion in *ompL* in all sequenced HE strains (**Supplementary File S2**). On top of the IS186B insertion, we found nine additional mutations in our parental DH10B strain compared to the reference Genbank sequence (CP000948.1) (**Supplementary File S2**). Intragenic mutations made up 82-84 % of the mutations in HE strains and 12-13 % of the intragenic SNPs caused truncation or frameshift mutations (**Supplementary Table S4**). We constructed HE isolate mutational spectra to analyze the frequency of the twelve possible SNPs and to compare these mutations to the spectrum induced by the mutagenic plasmid MP6 as observed by Badran et al^12^. (**Supplementary Fig. S5**). Our mutational spectra come from a much smaller number of mutations than in the Badran et al. study (hundreds vs. over one hundred thousand) and the conditions of the experiment are different, but in both cases MP6 appears to generate broad mutational spectra.

### Heat-evolved strains have mutations in essential genes, heat shock genes and other important genes

We analyzed genes mutated in ORFs and genes adjacent to intergenic mutations in our parental DH10B strain and in HE isolates. Intergenic mutations can alter the properties of promoters, operators, transcriptional terminators and ribosome binding sites. Additionally, intergenic mutations and mutations early in ORFs can also affect the translation initiation rates of the genes through changes in mRNA folding near the ribosome binding sites. Using the *E. coli* Keio collection of single gene knockouts of nonessential genes^14,15^, the TraDIS essential gene study^16^ and a σ^32^ regulon investigation^17^, we identified mutations in and around essential genes (**Table 1** and **Supplementary Table S5**). We did a similar analysis for heat shock genes (**Table 1** and **Supplementary Table S6**). For the missense mutations in essential and heat shock genes, we used DDGun^18^ to predict any changes in stability in the encoded proteins using published or Alphafold^19^ predicted structures and plotted the ΔΔG values (**Supplementary Fig. S6**). The essential proteins had an average ΔΔG of -0.59 and heat shock genes -0.29, indicating that on average, mutations in both sets of proteins were destabilizing. We found no nonsense mutations in essential nor heat shock genes. We found a few intragenic essential and heat shock gene mutations in genes located in the repeat region of the chromosome, and we have listed these mutations in the appropriate tables (**Table 1** and **Supplementary Table S6**). Mutations that decrease or abolish the function of the encoded proteins may have little meaning since the mutation frequency suggests that a wild-type copy of the mutated gene is still present in the chromosome, however, gain of function mutations are also possible. We also identified mutations in the strains that were lost, either through reversion, genetic changes in cultures grown for genomic DNA preparation or sequencing issues (**Supplementary Tables S7 and S8**).

**Table 1.**
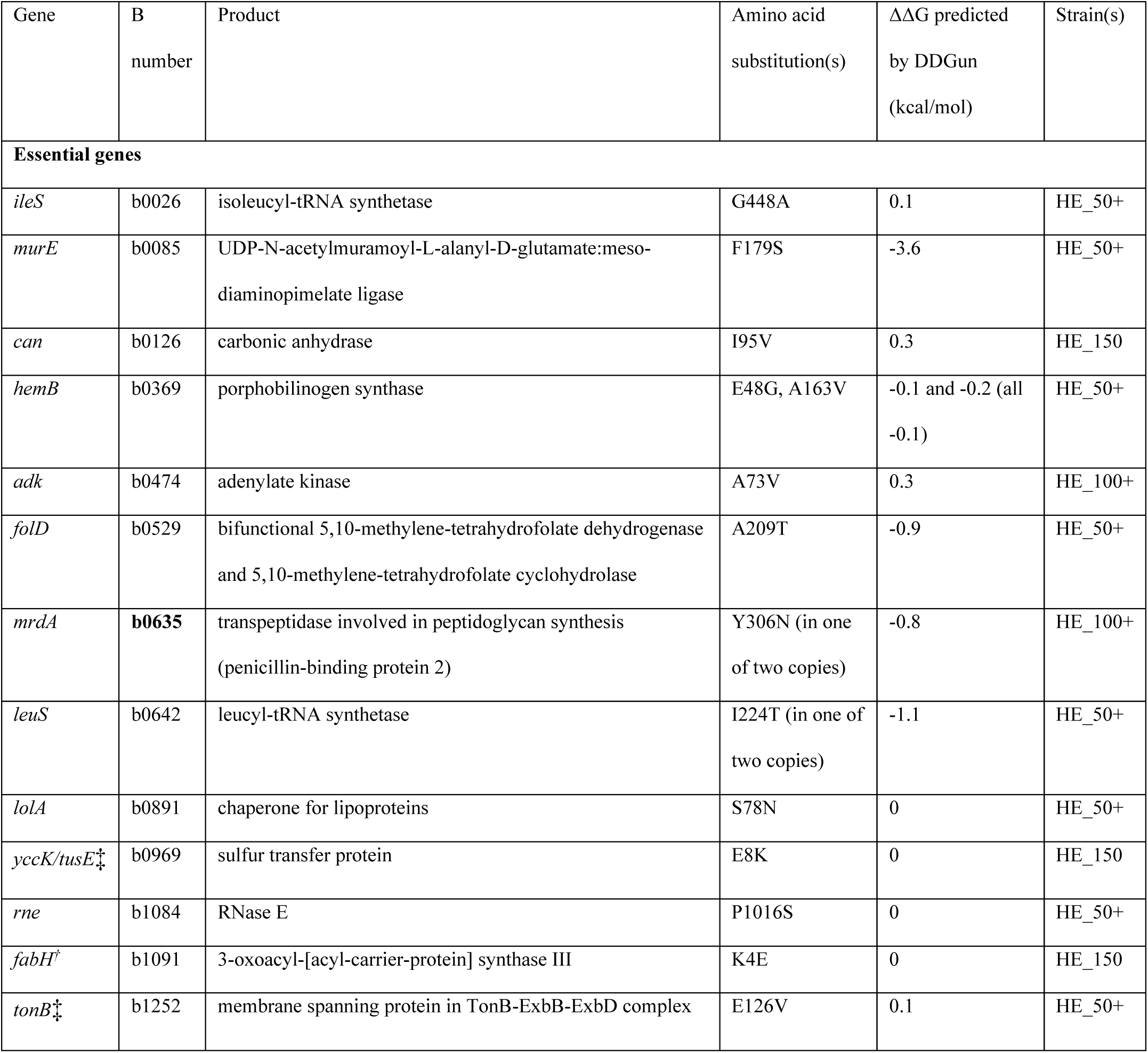
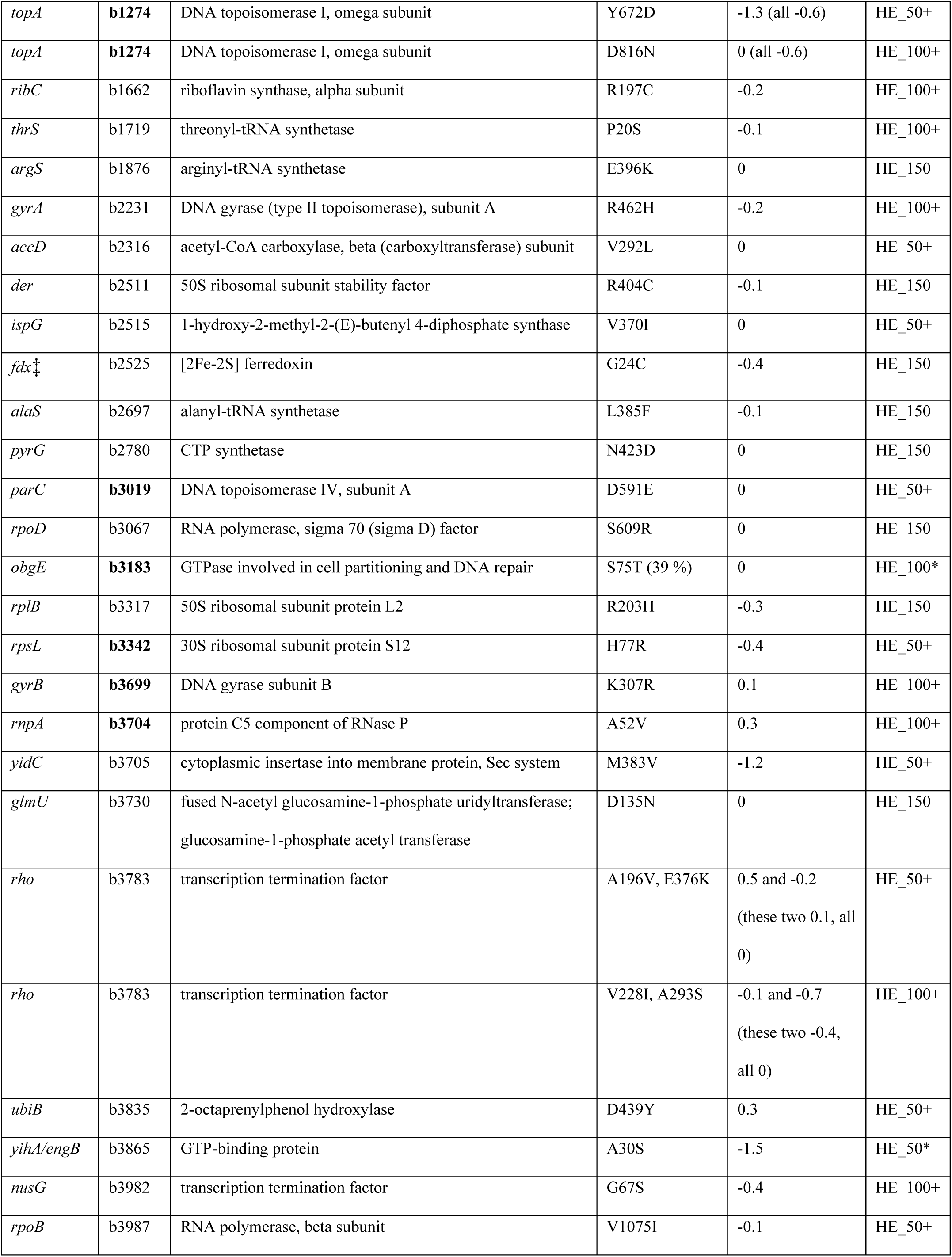
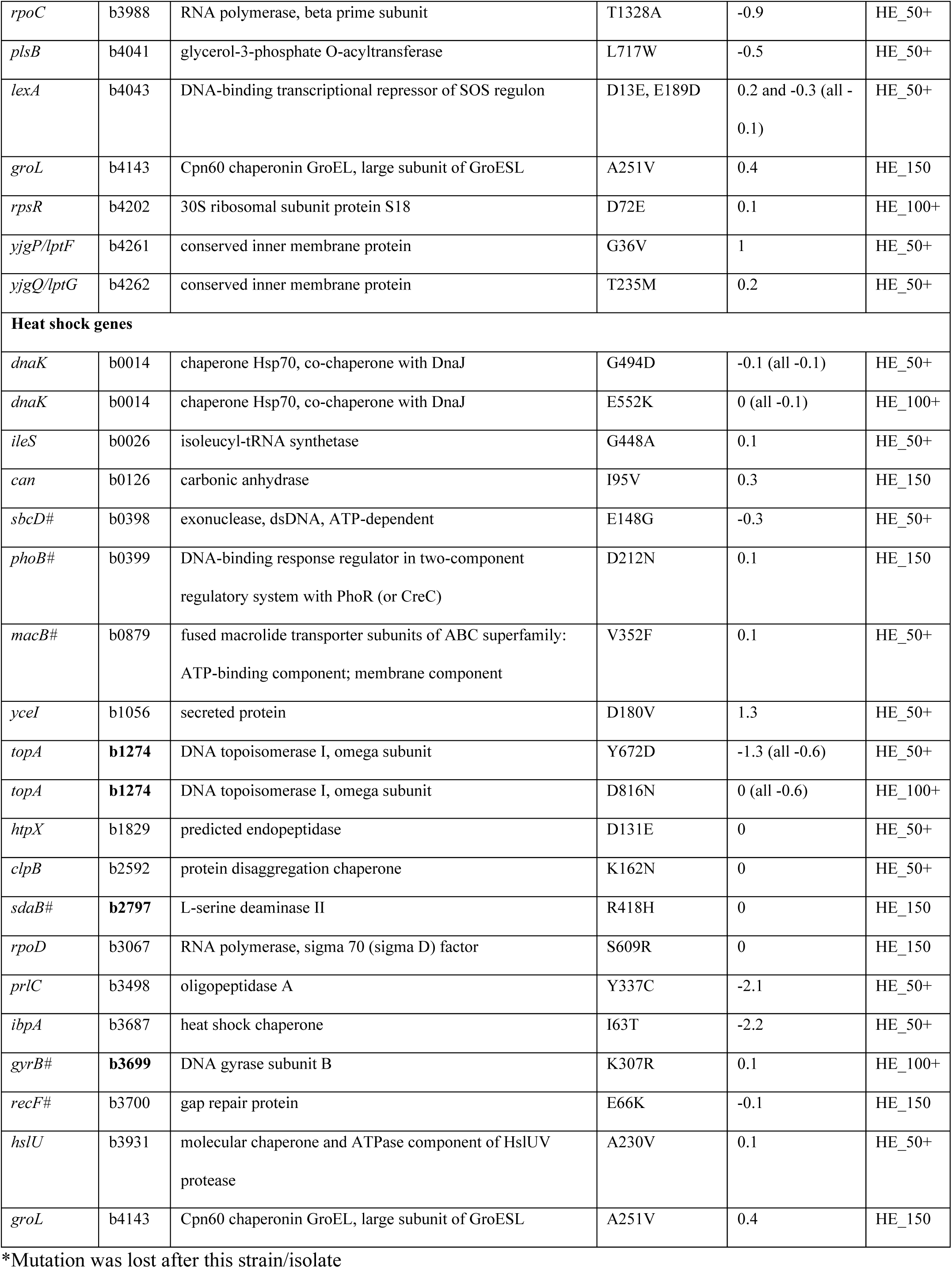
Missense mutations in essential and heat shock genes. For mutations at <70 % frequency, the percent frequency is shown in brackets in the Amino acid substitution(s) column. The b number is shown in bold if the melting temperature of the protein^33^ is below 52 °C (≤5 °C above the maximum LB broth growth temperature of HE_150). The DDGun^18^ predicted change in the free energy of unfolding between the wild-type and mutant proteins is shown, where a positive value indicates an increase in stability for the mutant protein and a negative value indicates a decrease in stability for the mutant protein. Where multiple amino acid substitutions are found in the same protein, the effects of the single mutations are shown first and in brackets the effects of combinations of mutations are shown.

We identified a few silent and intergenic mutations that may play important roles in adaptation to heat. For mutations near the start codons of genes with known transcription start sites, we calculated the predicted translation initiation rates^20^ for the genes with and without the mutation (**Supplementary Table S9**). One likely important change is a silent mutation in codon six of the essential heat shock gene *groS* in HE_150, which increases the predicted translation initiation rate^20^ from 17,753 to 24,547 au (**Supplementary Table S9**). On the transcriptional level, a SNP in the σ^32^ promoter of the heat shock gene operon *hslVU* brings the promoter closer to the σ^32^ consensus sequence^17^, and a SNP in the σ^32^ promoter of the heat shock gene operon *dnaKJ* shifts the promoter further away from the σ^32^ consensus sequence^17^. For a list of all parental DH10B and HE isolate mutations, see **Supplementary File S2.**

### Parental DH10B and HE_150 have different proteomes at both 37 °C and 42.5 °C

We wanted to determine if the genomic changes that we observed generated protein expression correlates, and thus we obtained label-free LC-MS/MS data from The University of Victoria Genome BC Proteomics Centre. Our samples included parental DH10B and HE_150 each grown at 37 °C and 42.5 °C, performed in duplicate. In each sample, 1,010-1,139 *E. coli* proteins were detected (**Supplementary File S3**). Among the proteins detected in all four samples, we compared protein abundances across the different strains and growth temperatures to look for general trends.

In previous studies where *E. coli* were adapted to a constant stressful temperature of 42 or 42.2 °C, they performed RNA sequencing analysis on the wildtype and adapted strains at 37 °C and 42-42.2 °C^8,9,11^. In those studies, they classified changes in gene expression at the transcript level into restored, unrestored, reinforced or novel expression patterns^9,11^. Restored protein expression indicates that there is a difference in protein abundance between DH10B grown at the two temperatures and that there is a difference between HE_150 and DH10B at 42.5 °C in the opposite direction. Unrestored refers to a difference in abundance between DH10B grown at the two temperatures but no difference in protein abundance between HE_150 and DH10B at 42.5 °C. Reinforced indicates a change in abundance, all in the same direction, of the following three comparisons: DH10B at 42.5 °C vs. 37 °C, HE_150 vs. DH10B at 42.5 °C and HE_150 at 42.5 °C vs. DH10B at 37 °C. Finally, a novel expression pattern means there is no difference between DH10B at both temperatures but there are differences in the following two comparisons, in the same direction: HE_150 vs. DH10B at 42.5 °C and HE_150 at 42.5 °C vs. DH10B at 37 °C. In the RNA sequencing data from studies performed at a constant stressful temperature, they found that the majority of transcripts had restored expression^8,9,11^. We wondered if we would observe that phenomenon at the protein level. We found 173 proteins with one of the four described expression patterns, including 30 restored (17.3 %), 32 unrestored (18.5 %), 9 reinforced (5.2 %) and 102 novel (59.0 %, **Supplementary File S3**). Thus, in HE_150, a novel expression shift is the most common expression shift observed.

In all expression categories except restored, there were significant overrepresentations in specific Gene Ontology biological processes, as well as a significant underrepresentation in one case (**Supplementary Tables S10** and **S11**). The unrestored category was enriched in proteins involved in protein folding (GO:0006457). The reinforced category had an overrepresentation of proteins involved in Mo-molybdopterin cofactor biosynthetic process (GO:0006777). Finally, the novel category was enriched in proteins involved in glycolytic process (GO:0006096), carboxylic acid catabolic process (GO:0046395), amino acid metabolic process (GO:0006520) and phosphate-containing compound metabolic process (GO:0006796), and depleted in proteins involved in regulation of cellular process (GO:0050794).

When we compared protein abundances in the strains close to their optimum growth temperatures (HE_150 at 42.5°C and DH10B at 37 °C), there were noteworthy differences for 218 proteins in both replicates, with 133 proteins at lower abundance in HE_150 at 42.5 °C than in DH10B at 37 °C and 85 proteins at higher abundance in HE_150 at 42.5 °C than in DH10B at 37 °C (**Supplementary File S3**). Of these 218 proteins, 156 had no intragenic changes nor intergenic changes between their gene and the genes immediately upstream and downstream. Proteins in higher abundance in DH10B were enriched in proteins involved in amino acid processes (aspartate metabolic process [GO:0006531], amino acid catabolic process [GO:0009063], alpha-amino acid biosynthetic process [GO:1901607]), pyruvate metabolic process (GO:0006090), carbohydrate catabolic process (GO:0016052), monosaccharide metabolic process (GO:0005996), protein homotetramerization (GO:0051289) and response to oxidative stress (GO:0006979), and depleted in proteins involved in regulation of cellular process (GO:0050794) and nucleic acid metabolic process (GO:0090304, **Supplementary Tables S12** and **S13**). Proteins in higher abundance in HE_150 were enriched in proteins involved in regulation of protein catabolic process (GO:0042176), histidine biosynthetic process (GO:0000105), Mo-molybdopterin cofactor biosynthetic process (GO:0006777), protein folding (GO:0006457) and response to heat (GO:0009408) and depleted in proteins involved in transmembrane transport (GO:0055085, **Supplementary Tables S12** and **S13**). Some of these results, particularly the depletion of proteins involved in membrane transport, may be influenced by the fact that our cell lysis method is optimized for cytosolic proteins and not optimal for isolating membrane proteins.

Zeroing in on particularly interesting mutated essential and heat shock proteins, DnaK, ClpB and HtpG had an unrestored expression and GroS had a reinforced expression. HslV, HslU, HtpG, GroL, GroS and ClpB had a higher abundance in HE_150 grown at 42.5 °C than DH10B grown at 37 °C. TopA was at a lower abundance in HE_150 at 42.5 °C compared to DH10B at 42.5 °C. We also looked at abundances of proteins that had mutations near the start codons of their genes for which we calculated predicted translation initiation rates (**Supplementary Table S9**). We detected noteworthy changes in protein abundances between HE_150 and parental DH10B for four of these proteins (SucA, FabH, RibB and GroS), and for three of the proteins (SucA, FabH and GroS) the change in abundance was shifted in the same direction as predicted by the calculated translation initiation rates (**Supplementary File S3** and **Supplementary Table S9**).

## DISCUSSION

In this work, we used a mutator plasmid to enhance mutagenesis and accelerate the ALE of *E. coli* to growth at higher temperatures. After 150 passages at elevated temperatures, we isolated a strain that was able to grow about 2°C above the maximal growth temperature of the parental DH10B strain (**Fig. 2**). This increase in the maximum growth temperature was achieved in five months in a strain with 1,059 mutations, whereas other comparable studies that did not employ a hyper-mutagenesis method raised the maximum growth temperature by 3 °C in a strain with ∼240 mutations^7^ in 6.9 years^2^, and 2 °C in a strain with 31 mutations in 8 months^4^. The HE_150 strain grew less well than the parental strain at moderate temperatures (**Fig. 3**) but was able to grow in the presence of levels of ethanol that completely inhibited the parental strain (**Supplementary Fig. S3**). We observed an average accumulation of 7-11 mutations per 24 hours, resulting in hundreds of genetic changes in each isolate (**Supplementary Table S4**). We found changes in heat shock genes encoding GroES, GroEL, DnaK, ClpB, IbpA and HslU (**Table 1** and **Supplementary Table S6**); changes in master regulator genes encoding components of the RNA polymerase (RpoB, RpoC and RpoD) and the transcriptional terminator Rho (**Table 1**); and changes that could affect the topology of DNA, and therefore potentially affect the expression level of several genes (**Table 1**). We believe that these mutations are likely the most important contributors to the thermotolerance of the strain, and they alone and in combination with the other mutations represent a unique set of mutations that have not been previously observed.

While it is difficult to ascribe the contribution to the heat-resistant phenotype of each of the hundreds of mutations that WGS uncovered, some themes are consistent with theoretical adaptation to heat and with results from previous adaptation studies^2–7^. These themes include a change in the expression of protein chaperones, mutations that affect the fluidity of cell membranes, genomic changes that affect DNA supercoiling, and missense mutations in transcription-association proteins. The latter two types of changes likely affect the expression of thousands of genes^11,21^. Indeed, our proteomic analysis of parental DH10B and HE_150 revealed important differences in protein abundances between the strains. When we compared the proteomes of the strains when grown around their optimum temperatures (DH10B at 37 °C and HE_150 at 42.5 °C) 218 proteins were at different abundances (**Supplementary File S3**).

We also identified proteins with restored, unrestored, reinforced and novel expression patterns, and found that the majority of the proteins had a novel expression pattern (59.0 %). In contrast, in RNA sequencing data from *E. coli* strains incubated at constant high temperatures (42.0 or 42.2 °C), the majority of the transcripts display a restored expression pattern^8,9,11^. We believe there are several factors that contribute to this difference. Due to different transcription, translation, transcript degradation and protein degradation rates, the correlation between transcript abundance and protein abundance varies for each protein. As well, the experimental conditions were quite different between our study and the studies performed at constant temperature, in terms of increasing vs. constant temperature, different starting strains, different media and different mutation rates (hyper-mutagenesis vs. spontaneous mutation). In the constant temperature studies the populations were cultured in minimal media, which leads to a lower maximum growth temperature due to the thermosensitivity of the enzyme MetA, as well as an increased number of genes that are essential under minimal media conditions. As well, in the constant temperature studies where hyper-mutagenesis is not employed, the growth environment is kept constant, and investigators found that initial beneficial mutations lead to a higher fitness boost and changes in expression for more genes than later changes, which often compensate for detrimental expression changes in a smaller number of genes^8,9,11^. We believe that in a changing environment, there is no set of beneficial mutations that are indefinitely helpful in the environment due to its changing nature. We also expect that with hyper-mutagenesis, more mutations which are detrimental hitchhike alongside beneficial and neutral ones, which offer another level of challenges for the cells to adapt to. Since the final incubation temperature in our experiment exceeds the maximum growth temperature of the starting strain, restorative shifts in gene expression are likely not sufficient for allowing growth above that temperature. Mutations which directly or indirectly facilitate the activity of essential processes that normally require a thermosensitive protein are likely also necessary, which could in some cases be accomplished by a novel expression pattern.

Mutations in heat shock genes were a prominent aspect of genomic changes that we observed. The silent mutation in codon six of the essential heat shock gene *groS* in HE_150 increases the predicted translation initiation rate of *groS*, and, through translational coupling^22^, it could also increase the translation of the *groL* gene downstream. Proteomic analysis revealed different abundances in GroS and GroL in HE_150 compared to parental DH10B (**Supplementary File S3**). At 37 °C, GroS levels are 2.8-3.2 X times higher and GroL levels 1.8-2.7 X times higher in HE_150 compared to parental DH10B. At 42.5 °C, GroS levels are 2.0-2.6 X times higher and GroL levels 1.6-2.2 X times higher in HE_150 than in parental DH10B. HE_150 additionally has an A251V substitution in the apical domain of GroEL. The effect of this change is unknown, but it is predicted to be stabilizing by DDGun (**Tables 1** and **2**).

Previous work clearly demonstrated that increases in GroESL expression enhance *E. coli*’s ability to grow at higher temperatures^2^. We recently analyzed the strain developed in that study and demonstrated that the hyper-expression of GroESL was due to the presence of a plasmid-borne copy of the *groESL* operon and that the presence of this recombinant plasmid was sufficient to raise the thermotolerance of *E. coli* strains^7^.

It is clear how a molecular chaperone that is overexpressed during heat shock could be beneficial for high-temperature growth, and there are also interesting studies looking at the relationship between mutations and molecular chaperone overexpression^23–25^. Various chaperones have been shown to either buffer or potentiate the effects of amino acid substitutions in proteins^26,27^. For example, a chaperone could assist a protein in reaching its native fold despite the mutation, but without chaperone access the protein misfolds or aggregates, leading to phenotypic differences under different conditions. Some studies highlight links between mutation rates and chaperone overexpression, where chaperone expression increases as cells accumulate mutations^23^ and where proteins that are clients of molecular chaperones are able to evolve more quickly (i.e., acquire more mutations per unit time) than proteins that fold independently^24,28^. Therefore, one role of higher expression of GroESL in HE cells may be to allow the cell to tolerate the large number of missense mutations affecting expressed proteins.

The *hslVU* operon encodes a heat shock protease complex that degrades misfolded proteins. The mutation upstream of *hslVU* (also known as *clpQY*) may drive higher expression of these genes under heat shock conditions, as it brings the promoter sequence closer to the σ^32^ consensus sequence (**Supplementary Table S6**). In support of this, we did detect a higher abundance of HslU and HslV in HE_150 than DH10B both grown at 42.5 °C, though this only ranged from 1.6-2.1 X higher (**Supplementary File S3**). In HE cells, it could be that there is a shift towards more degradation and less refolding of unfolded or misfolded proteins. In HE_50+, HslU also has an A230V substitution in the I-domain that binds substrate proteins which is predicted to be stabilizing by DDGun (**Table 1**).

The promoter driving the heat shock operon *dnaKJ* has suffered a mutation in HE_50 and later strains that may decrease expression of the genes under heat shock conditions, as it brings this sequence further away from the σ^32^ consensus sequence (**Supplementary Table S6**). However, *dnaKJ* has two other σ^32^ promoters in between the mutated one and the start of *dnaK*, so it is not clear how much of an effect this mutation has on the operon.

Consistent with this mutation, there is an increase in DnaK abundance from 37 °C to 42.5 °C in parental DH10B (2.3-2.6 X) but not in HE_150 (1.8 X, **Supplementary File S3**). As well, at 42.5 °C, the DnaK abundance is lower in HE_150 than DH10B (0.5-0.7 X, **Supplementary File S3**). In HE_50+, the strains also have a G494D mutation in *dnaK* predicted to be destabilizing by DDGun (**Table 1**). Work by General et al.^29^ predicts that Gly494 is a sensor residue, meaning that it senses allosteric signals from one or more binding partners of DnaK (DnaJ, ATP, ADP, substrate protein, etc.). Thus, a substitution at this residue could alter the function of the DnaK/DnaJ/GrpE chaperone system in HE cells. In HE_100+, the strains have an additional amino acid substitution in DnaK; E552K which is predicted to be neither stabilizing nor destabilizing by DDGun (**Table 1**). Both G494 and E552 reside in the substrate binding domain (SBD) of DnaK, with G494 present in the beta part of the SBD and E552 present in the alpha lid domain of the SBD.

In adapting to higher incubation temperatures, HE strains gained increased resistance to ethanol (**Supplementary Fig. S3**). Organic chemicals like ethanol mimic a heat shock in many ways, through envelope stress and protein unfolding. Perhaps owing to this, thermophiles are more resistant to organic solvents than mesophiles or psychrophiles. Our results are consistent with previous studies^30,31^ and support the notion that adapting to high temperatures goes hand in hand with increased ethanol resistance.

Other important changes in HE strains are mutations in master regulators (**Table 1**). In HE_50+, the isolates have mutations in components of the RNA polymerase RpoB (V1075I) and RpoC (T1328A). Between HE_100 and HE_150, a mutation in the housekeeping sigma factor σ^70^ (RpoD) emerged (S609R). An RNA polymerase binding partner which terminates transcription for ∼20 % of *E. coli* genes^32^, Rho, also has two mutations in HE_50+ (A196V and E376K) and two additional mutations in HE_100+ (V228I and A293S). In HE_100+, cells have a G67S mutation in the transcription termination/antitermination factor NusG. HE_150 also has the substitution D90E in the RNA polymerase binding transcription factor DksA (**Supplementary File S2**). Interestingly, the heat-evolved *E. coli* strain BM28^2,7^ has a D90N mutation in DksA suggesting that changes to D90 could be an adaptation to heat.

Finally, HE isolates acquired mutations in five ribosomal protein genes and two rRNA genes (**Supplemental Table S6** and **Supplementary File S2**). These changes almost certainly cause changes in gene expression in HE cells^8–11^. Indeed, amongst the 218 proteins with noteworthy differences in abundances between DH10B at 37 °C and HE_150 at 42.5 °C, 156 (71.6 %) had no intragenic mutations nor were they immediately adjacent to any intergenic mutations (**Supplementary File S3**). For some of these proteins, these mutations in master regulators are likely the cause of their change in expression level.

In this work, we found genomic changes that are commonly observed in ALE studies, including changes in RNA polymerase components, Rho and TopA. We also found mutations that have been previously observed specifically in studies of ALE to heat, namely GroESL^2,7^, GlpF^4^, CytR^3,7^, and Gpp^3,7^. Through WGS, we detailed a unique evolutionary route towards growth at high temperature in *E. coli*. After five months of ALE, the final strain achieved a maximum plate growth temperature ∼2 °C higher than the parental DH10B strain. As well, we found many differences in protein abundance in HE_150 and the starting DH10B strain at both 37 °C and 42.5 °C, including for important heat shock proteins DnaK, ClpB, GroS, GroL, HslV and HslU. We hope that this study contributes to a collective understanding of evolution to greater heat tolerance in *Escherichia coli*, and a better understanding of the general mechanisms of evolution towards increased thermotolerance in other organisms.

## METHODS

### ALE to heat with MP6 hyper-mutagenesis

Chemically competent DH10B was transformed with plasmid MP6 and plated on LB agar (10 g/L tryptone, 5 g/L yeast extract, 5 g/L NaCl and 15 g/L agar) containing 25 µg/mL chloramphenicol and 0.1 w/v % glucose. For ALE experiments, cells were cultured for 24-hour periods in 2 mL of 2xYT media (16 g/L tryptone, 10 g/L yeast extract and 5 g/L NaCl) containing 25 µg/mL chloramphenicol and 0.1 mM arabinose with shaking at 100 rpm. Cells were serially cultured by diluting 1:100 or 1:1000 every 24 hours into fresh media, usually at 1:1000 and exclusively at 1:1000 in the latter half of the ALE experiment. Occasionally, mostly near the beginning of the evolution, the cells were streaked onto LB agar plates with chloramphenicol and 0.1 % glucose for 72 hours and incubated at 46.0-47.3°C. Isolated colonies from these plates were used to start up fresh liquid cultures, and these plate incubations were not considered as adaptation steps. The temperature of the incubator for the heat ALE experiments was slowly increased over 150 days, from 43.5 °C to 46.9 °C. Incubator maximum, minimum and average temperatures were measured with a Fluke 53IIB digital thermometer equipped with a K-type thermocouple and recorded over 24-hour periods, and the temperatures reported represent average temperatures over the 24-hour period. All temperatures reported should be considered to have a potential error of ± 0.2 °C.

Periodically (at HE_50, HE_80, HE_100, HE_110, HE_123 and HE_150), HE cells were cured of their MP6 plasmids and retransformed with parental MP6. To generate a plasmid-free isolate, a single colony from a high temperature (46.5-47.3 °C) plate was used to inoculate LB with 0.1 % or 1 % w/v SDS, and cells were cultured in SDS media every 24 hours for 72 hours at 37 °C. A dilution of the culture was plated on LB agar and 30 colonies were streaked onto LB and LB + chloramphenicol agar to isolate HE strains cured of MP6. MP6-cured HE strains were then transformed with fresh MP6 and ALE to heat was continued. Strains HE_50, HE_100 and HE_150 were derived from single colonies cured of MP6 from the ALE populations; thus, they represent a single individual from the ALE populations at each time point.

### Whole-genome sequencing of parental DH10B and HE strains

20 mL LB cultures of our laboratory strain of DH10B (“parental DH10B”) and HE cells were incubated at 200-250 rpm overnight at 37 °C and 42 °C, respectively. Cells were harvested from the cultures, suspended in TE, and lysis buffer components were added and mixed into the cell suspension by inverting the tubes. The resulting lysis buffer was composed of 0.5 % w/v SDS, 100 µg/mL Proteinase K, 100 µg/mL RNAse A and 10 mM CaCl_2_ in TE buffer. Lysis was carried out for 2 hours at 37 °C with mixing by inversion every 30 minutes or for 15 minutes at 55 °C. The gDNA was extracted twice with one volume (1 mL) of 25:24:1 phenol:chloroform:isoamyl alcohol, and the resulting aqueous layer was purified via spin column. gDNA samples were sent to the Microbial Genome Sequencing Center (Pittsburgh) for Illumina sequencing (150 Mb, ∼32 X coverage for DH10B-related cells) and Oxford Nanopore sequencing (300 Mb, ∼64 X coverage for DH10B-related cells).

For Illumina data, FastQ paired-end read files were imported into Geneious Prime 2022 using the default settings with the insert size set to 500 bp. Reads were trimmed using BBDuk with the default settings and trim adapters selected (“trim low quality” set to both ends, “minimum quality” of 30, “trim adapters based on paired read overhangs” set to a minimum overlap of 24 and “discard short reads” set to a minimum length of 30). We also merged the pair-end reads with BBMerge^33^ in order to generate longer reads where possible (when the paired-end reads overlapped). For Nanopore data, FastQ files were imported into Geneious Prime, and Nanopore was selected as the data type.

### Assembly of the parental DH10B and HE isolate genomes

The parental DH10B and HE genomes were assembled using a combination of the SPAdes *de novo* assembly algorithm, the Geneious Prime map to reference function and manual curation/editing. Trimmed Q30 (minimum quality 30) Illumina reads and raw Oxford Nanopore reads from HE_150 gDNA were used for SPAdes *de novo* assembly. Three large HE_150 scaffolds were built which covered all but ∼120 kb of the DH10B genome. This missing ∼120 kb corresponds to part of the 226 kb repeat region of DH10B, which is a tandem duplication of 113 kb of the genome. Based on Illumina read coverage and Nanopore reads, this region is not missing and is instead likely an artifact of the difficulty of *de novo* assembly of repeat regions. In the three *de novo* scaffolds, some regions containing sequences repeated elsewhere in the genome including tRNA genes and REP repeat regions had small indels and/or runs of N nucleotides. When we mapped the trimmed or raw Illumina HE reads to the DH10B reference sequence (CP000948.1), and when we looked at the three HE_150 SPAdes scaffolds, there was only evidence for two changes larger than a few bp, in the form of additional insertion sequence elements. Therefore, the indels and/or runs of N’s in the tRNA and REP regions appeared to be assembly artifacts, and we deduced that there were no indels in those regions.

Using the contigs generated by the “map to reference” runs and the scaffolds generated by the SPAdes run, we built a HE draft genome composed of the DH10B reference sequence with the two added IS elements. When we mapped our parental DH10B trimmed or raw Illumina reads to the DH10B reference sequence and the HE draft genome sequence, we could see that only one of the two HE IS insertions were present in the parental DH10B genome. Thus, we built a DH10B draft genome with just one added IS compared to the reference DH10B genome. We mapped trimmed and raw HE Illumina reads to the HE draft genome and trimmed and raw parental DH10B Illumina reads to the parental DH10B draft genome. With those files, we checked to see that there was no evidence for any larger changes, and we used the consensus sequences from those maps to reference runs as draft genomes for parental DH10B and each HE isolate. The consensus sequences were determined using the setting “Highest Quality (60 %)”, where the nucleotide reported as the consensus at each position makes up at least 60 % of the reads adjusted for quality. For some mutations, about half of the reads contained the reference nucleotide and about half of the reads contained a single variant nucleotide. In this case, the consensus nucleotide was called as both/either nucleotide using the appropriate non-ATGC nucleotide code for the two nucleotides (ex: R = A or G). Finally, we manually verified these draft genomes by mapping the appropriate Illumina reads to them, looking for variations/SNPs and editing the draft genomes if need be to include any more high-frequency variations.

To identify variations compared to the DH10B reference sequence, excluding the IS element insertions, we mapped Illumina reads to the appropriate draft genome (the parental DH10B draft genome for parental DH10B and the HE draft genome for HE isolates) using the default settings of “map to reference” with “do not trim” selected. Using the contig generated by mapping the paired-end reads to the reference, we found variations using the default settings of “find variations/SNPs” and exported them for analysis in Microsoft Excel.

### Growth experiments

All growth experiments were performed with plasmid-free parental DH10B and HE isolates. For all growth experiments, strains were plated from glycerol stocks onto LB agar plates and incubated at 37 °C overnight. For high-temperature plate growth experiments, colonies were used to inoculate liquid LB, grown at 42 °C, normalized to 1 OD_600_ unit and frozen at -80 °C in 10 % glycerol. Stocks were thawed on ice, and 10 µL of 10^-2^, 10^-3^, 10^-4^, 10^-5^ and 10^-6^ dilutions were spotted onto LB agar plates in duplicate. Plates were imaged after 24 hours of incubation at each temperature (37, 45, 46.8 and 47 °C). For liquid growth experiments, several colonies from a 37 °C plate were used to inoculate 2 mL of LB or LB with 4 % v/v ethanol in 16 mm diameter glass test tubes, and the tubes were shaken at 250 rpm in quintuplicate. The 4 % v/v ethanol growth experiment was performed at 37.3 °C. For doubling time experiments, absorbances at 600 nm were read and recorded every 20 minutes, and five biological replicates were used per strain. For each strain, optical densities ≤0.4 excluding time points 0 and 20 minutes were used to calculate the doubling times^34^ using GraphPad Prism 9’s exponential growth equation. Error bars were calculated by GraphPad Prism 9 and represent standard deviations from the mean, except for the doubling time graph where they indicate the 95 % confidence interval.

### Proteomic analysis of parental DH10B and HE_150

Parental DH10B and HE_150 were streaked onto LB agar and incubated at 37 °C overnight. Colonies were scraped from the plates, suspended in LB broth and diluted to an absorbance at 600 nm of 0.1 in 5 mL LB broth. The cultures were grown at 37 °C and 42.5 °C for 3.5 hours, the final absorbances at 600 nm were recorded and 4 mL of each culture was pelleted and the LB media was discarded. The cells were resuspended in 0.5 mL of PBS buffer with 0.3 g of acid-washed glass beads. The cells were vortexed for 30 seconds and incubated on ice for 30 seconds, and this was repeated three additional times. The lysis and bead mixture was centrifuged at 21,000 rpm for 4 minutes. Approximately 0.4 mL of supernatant was moved to a new tube, and the protein concentration was measured by Qubit. All protein concentrations were ∼1 mg/mL. Samples were split into 50 µL aliquots and frozen at -20 °C until used. Aliquots containing 20 µg of protein from two biological replicates grown and prepared at separate times were analyzed by label-free quantitative LC-MS/MS by The University of Victoria Genome BC Proteomics Center. See **Supplemental File S1** for proteomic method details.

Protein abundances were used to identify changes in gene expression as described previously^9,11^, including restored, unrestored, reinforced and novel expression patterns. Protein abundance ratios >2 and <0.5 were considered noteworthy. Restored protein expression indicates that there is a noteworthy difference in protein abundance between DH10B grown at 37 °C and 42.5 °C and that there is a noteworthy difference between HE_150 and DH10B at 42.5 °C in the opposite direction. An example of this would be a protein being at a higher abundance at 42.5 °C than 37 °C in DH10B and the protein being at a lower abundance in HE_150 at 42.5 °C compared to DH10B at 42.5°C. Unrestored refers to a noteworthy difference in abundance between DH10B grown at 37 °C and 42.5 °C but no noteworthy difference in protein abundance between HE_150 and DH10B at 42.5 °C. Reinforced indicates a noteworthy change in abundance, all in the same direction, of the following three comparisons: DH10B at 42.5 °C vs. 37 °C, HE_150 vs. DH10B at 42.5 °C and HE_150 at 42.5 °C vs. DH10B at 37 °C. Finally, a novel expression pattern means there is no noteworthy difference between DH10B at 42.5 °C and 37 °C but there are noteworthy differences in the following two comparisons, in the same direction: HE_150 vs. DH10B at 42.5 °C and HE_150 at 42.5 °C vs. DH10B at 37 °C.

## AUTHOR CONTRIBUTIONS STATEMENT

BM performed the experiments and prepared the manuscript, figures, tables and supplemental materials. FN performed an experiment and supervised. Both authors reviewed the manuscript.

## DATA AVAILABILITY

All data generated and analysed during this study are included in the manuscript, its Supplementary Information or are available from Genbank. Supplementary File S1 contains the Supplementary Figures and Tables, and Supplementary File S2 is an Excel file containing all of the mutations in parental DH10B and the HE strains. The assembled genomes are available through the accession numbers CP110014-CP110018 and the raw Illumina and Oxford Nanopore reads are available through the accession numbers SRR21983746-SRR21983750. All data can be accessed through the Genbank BioProject PRJNA892151 (https://www.ncbi.nlm.nih.gov/bioproject/PRJNA892151). HE isolates are available through Addgene: HE_50 (Addgene ID 201049), HE_100 (Addgene ID 201050) and HE_150 (Addgene ID 201051).

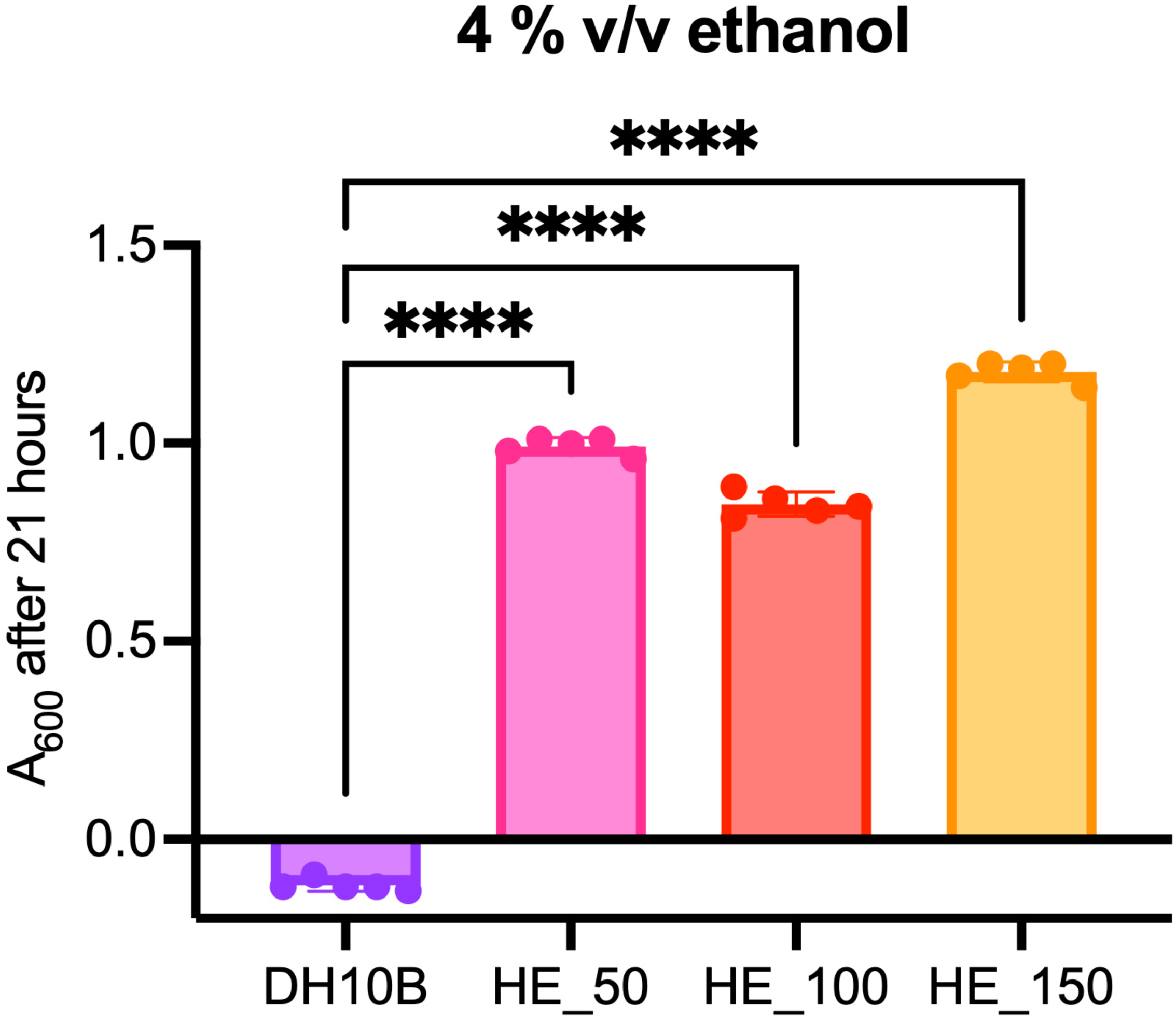

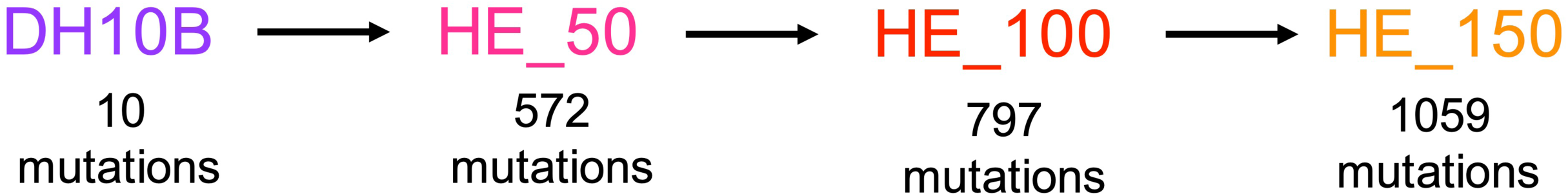

