## Supplementary File S1 for "Genomic analysis of laboratory-evolved, heat-adapted *Escherichia coli* strains"

Supplemental Material for **“**Genomic analysis of laboratory evolved, heat adapted *Escherichia coli* strains”

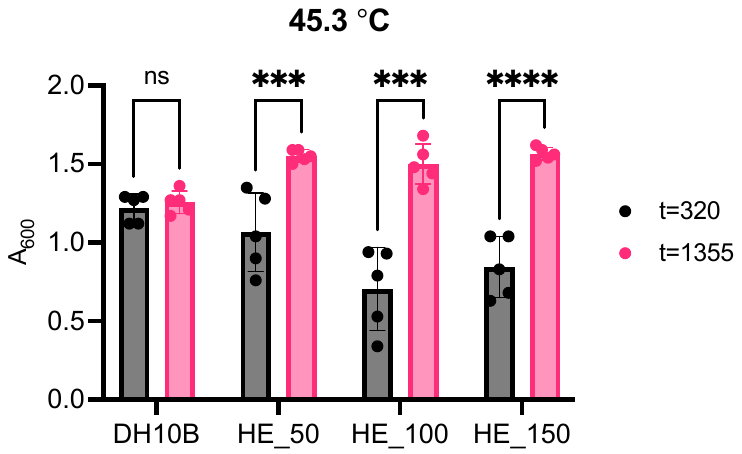

**Supplementary Figure S1.** Comparison of optical densities at 320 minute (5 hours 20 minutes) and 1355 minute (22 hours 35 minutes) timepoints for parental DH10B and HE isolates grown at 45.3 °C. Five replicates were used for each strain and the absorbance was measured at 600 nm. Comparisons are indicated with lines and ns indicates not significantly different, *** indicates a P value ≤0.001 and **** indicates a P value ≤0.0001.

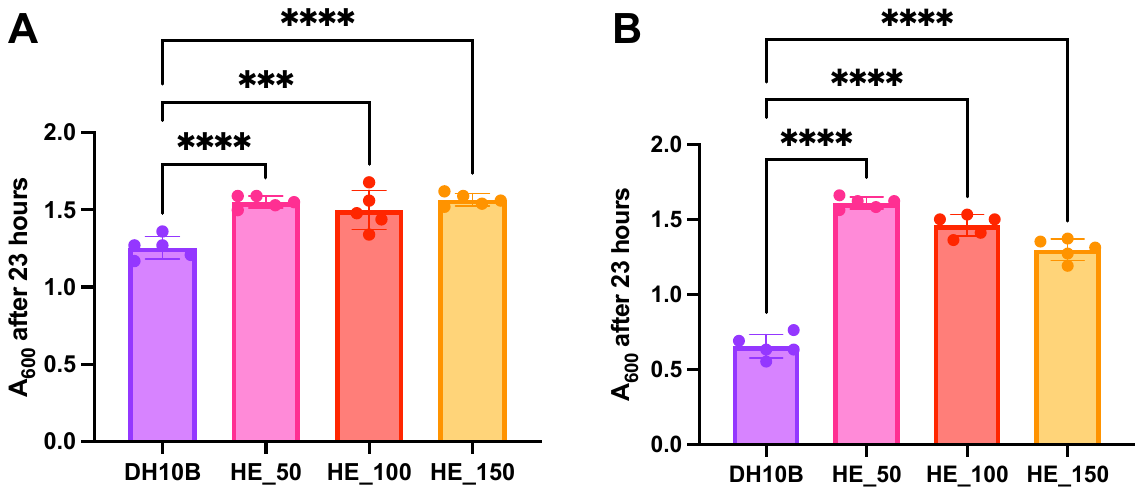

**Supplementary Figure S2.** Final optical densities of parental DH10B and HE cultures at 45.3 °C (**A**) and 46.3 °C (**B**). Five replicates were used for each strain and the absorbance was measured at 600 nm. Comparisons are indicated with lines and *** indicates a P value ≤0.001 and **** indicates a P value ≤0.0001.

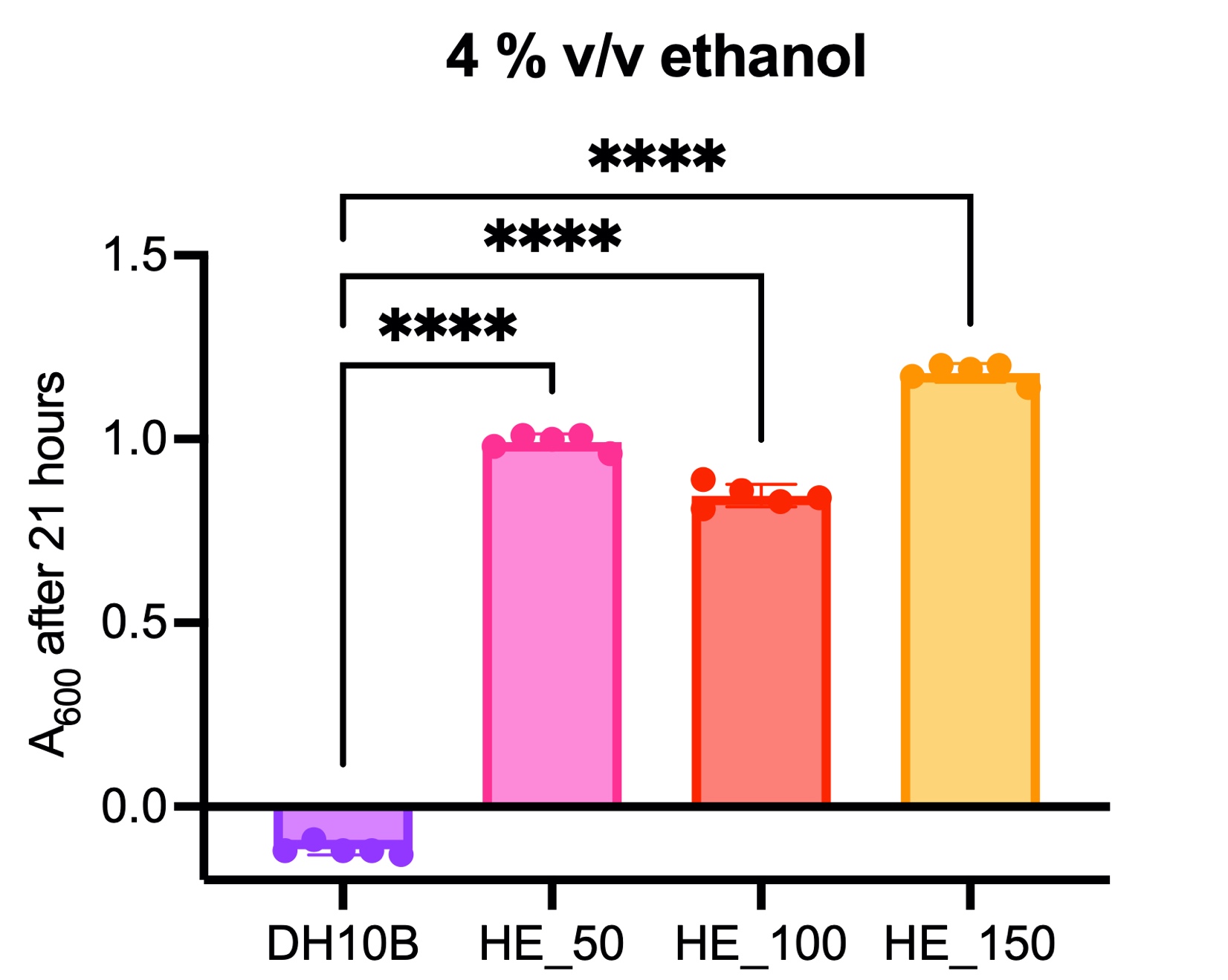

**Supplementary Figure S3.** HE isolates acquired ethanol resistance. Final optical densities of parental DH10B and HE cells grown in 4 % v/v ethanol in LB. Comparisons between parental DH10B and HE isolates are indicated with lines and **** indicates a P value ≤0.0001.

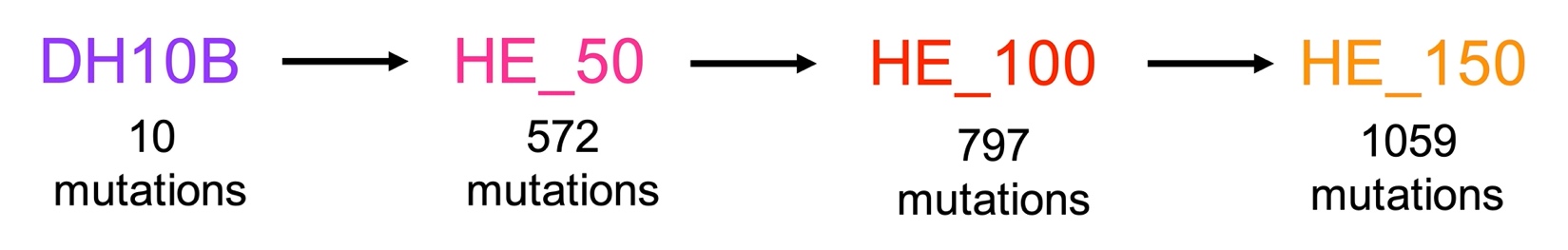

**Supplementary Figure S4.** Numbers of mutations in HE isolates and the number of mutations that arose (indicated by the “+” sign) or reverted (indicated by the “-” sign) between each isolate.

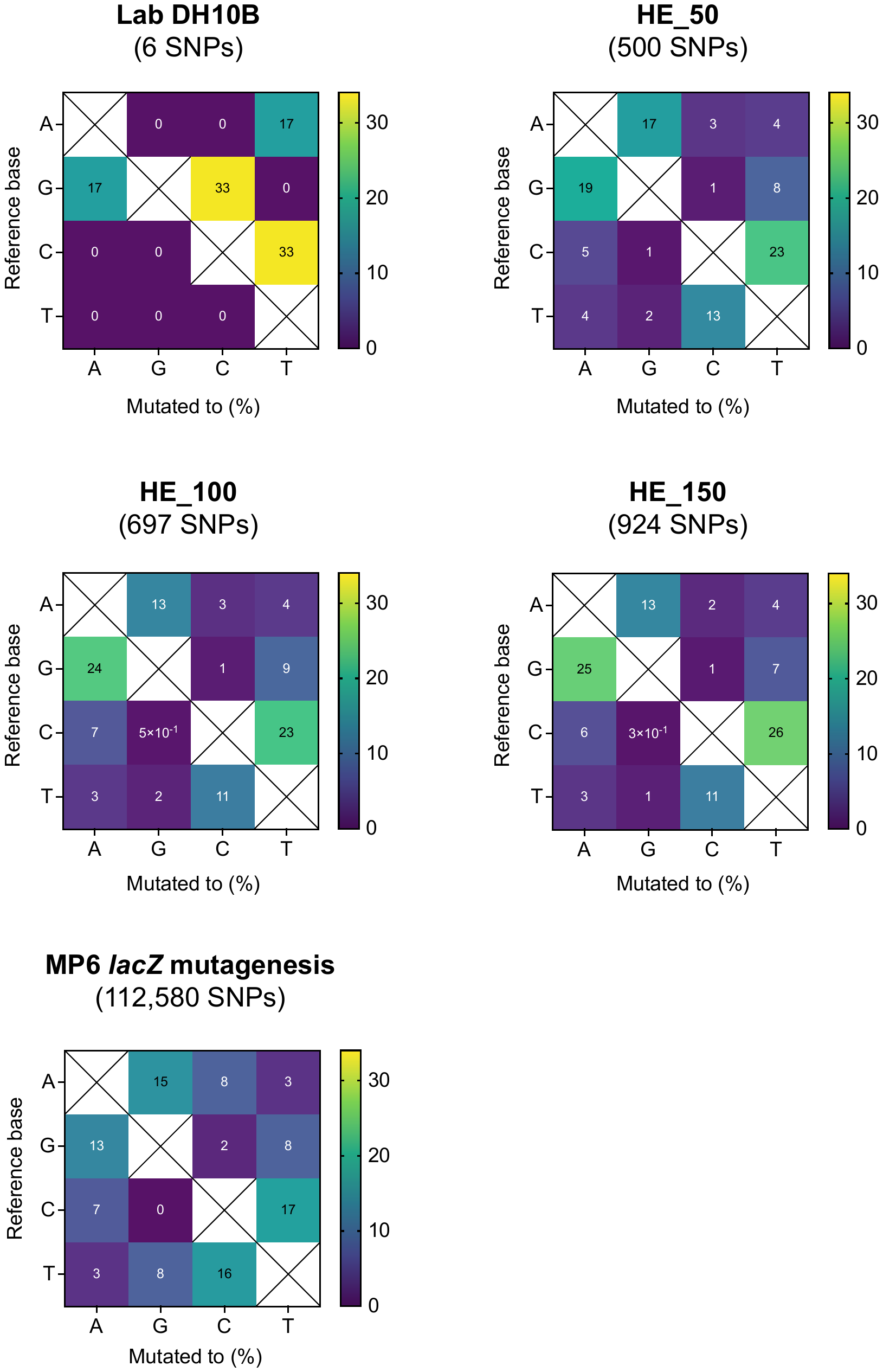

**Supplementary Figure S5.** Mutational spectra from MP6 mutagenesis in HE cells. The MP6 mutational spectrum determined by phage *lacZ* mutagenesis by Badran et al. (bottom right panel) has been recreated from their paper for comparison^12^.

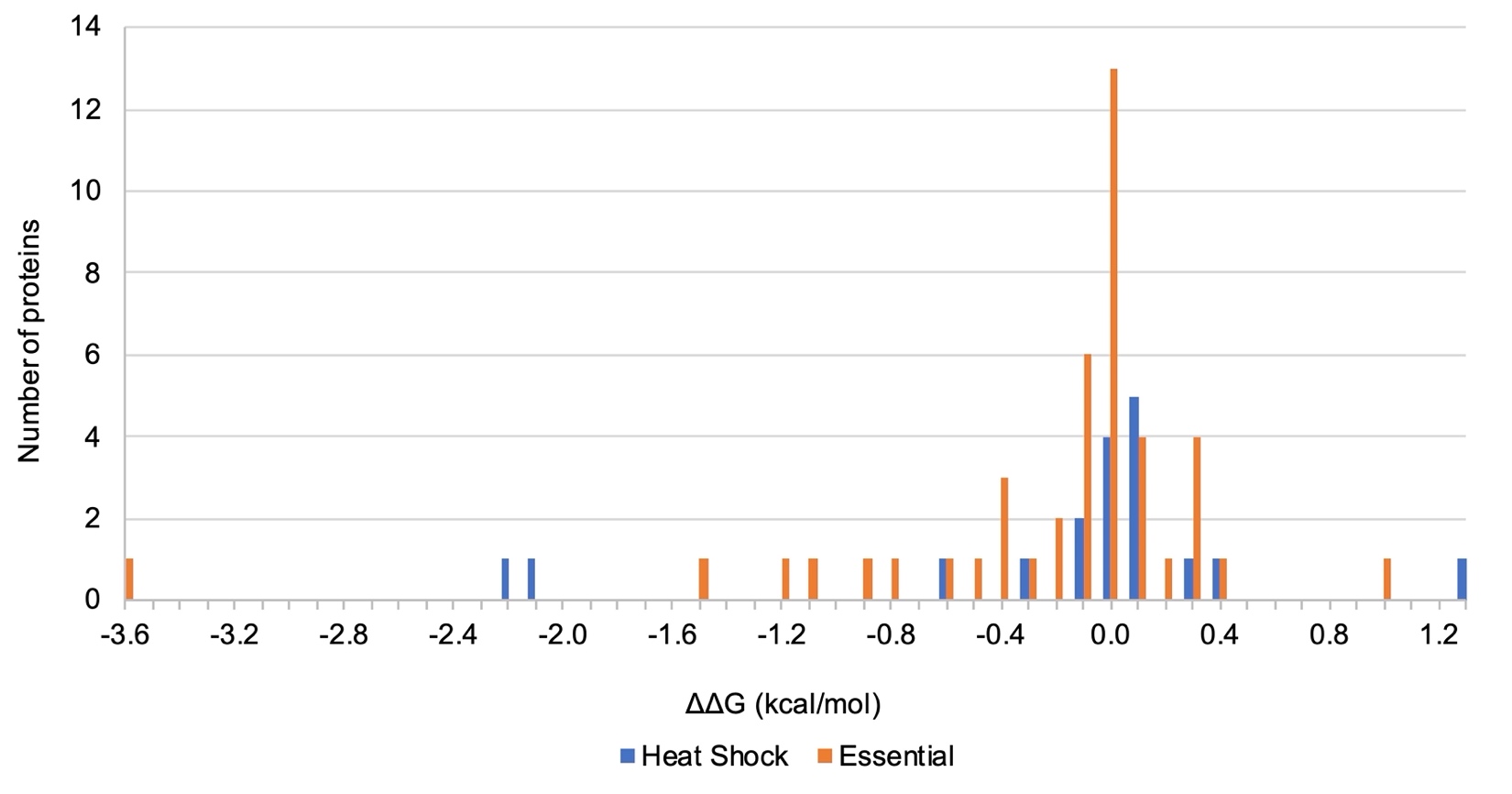

**Supplementary Figure S6.** Changes in protein stability in heat shock and essential proteins containing substitution mutations. The predicted change in the free energy of unfolding (ΔΔG) calculated by DDGun^18^ is shown, where positive values indicate an increase in stability and negative values indicate a decrease in stability.

**Supplementary Table S1**. NextSeq 2000 Illumina WGS data statistics.

| Strain | Reference genome | Number of trimmed paired-end reads mapped to reference | Mean coverage when mapped to reference genome | Standard deviation of coverage | Minimum coverage | Maximum coverage | Pairwise identity (%) | Mean read lengths | Mean pair distance |
| --- | --- | --- | --- | --- | --- | --- | --- | --- | --- |
| Parental DH10B | DH10B | 2063658 | 58 | 41.5 | 2 | 614 | 99.8 | 131.8 | 248 |
| HE_50 | DH10B | 2714300 | 74.2 | 34.3 | 3 | 506 | 99.8 | 128.3 | 230 |
| HE_100 | DH10B | 2833240 | 76.2 | 44.3 | 0 | 537 | 99.8 | 126.2 | 217 |
| HE_150 | DH10B | 2553658 | 67.8 | 27 | 1 | 393 | 99.8 | 124.7 | 214 |

**Supplementary Table S2.** HE_150 Oxford Nanopore WGS data statistics.

| Strain | Reference genome | Number of reads | Mean length | Standard deviation of length | Minimum length | Maximum length | Number of bases covered by reads | Mean coverage per base |
| --- | --- | --- | --- | --- | --- | --- | --- | --- |
| HE_150 | DH10B | 82131 | 5060.8 | 5418.6 | 48 | 64671 | 415650745 | 88.7 |

**Supplementary Table S3.** Number of repeat region mutations in parental DH10B and HE cells.

| Strain | Number of repeat region mutations | Expected number of mutations in a ~227 kb region |
| --- | --- | --- |
| Parental DH10B | 0 | 1 |
| HE_50 | 34 | 27 |
| HE_100 | 40 | 38 |
| HE_150 | 53 | 51 |

**Supplementary Table S4.** Parental DH10B and HE mutation statistics. For identifying changes including SNPs and small indels and substitutions, raw paired-end Illumina reads were trimmed with BBDuk2 and mapped to reference in Geneious Prime 2022 using the appropriate reference genome. Variations/SNPs were found with Geneious Prime 2022.

| Strain | Total changes (SNPs + small indels + IS insertions) | Average # of changes per 24 hours | Number of SNPs | Percent SNPs of all changes (%) | Percent intragenic mutations (%) | Percent intragenic SNPs causing truncations and frameshifts (%) |
| --- | --- | --- | --- | --- | --- | --- |
| Parental DH10B | 10 | N/A | 6 | 60 | 70 | 17 |
| HE_50 | 572 | 11.4 | 500 | 87 | 84 | 12 |
| HE_100 | 797 | 8 | 697 | 87 | 83 | 13 |
| HE_150 | 1059 | 7.1 | 924 | 87 | 82 | 12 |

**Supplementary Table S5.** Intergenic and silent mutations in or around essential genes. For intergenic mutations, the essential genes are indicated with asterisks, arrows surrounding genes indicate their directions and genes in bold have the insertion upstream of their start codons, meaning the insertions may affect the transcriptional and/or translational regulation of the genes. The b number is shown in bold if the melting temperature of the protein^33^ is below 52 °C (≤5 °C above the maximum LB broth growth temperature of HE_150). If a mutation is at <70 % frequency, the frequency is shown in brackets in the Mutation(s) column.

| Gene(s) | B number(s) | Product(s) | Mutation(s) | Strain(s) |
| --- | --- | --- | --- | --- |
| *rpsT*‡ | b0023 | 30S ribosomal subunit protein S20 | silent codon 61 | Parental DH10B+ |
| *murE* | b0085 | UDP-N-acetylmuramoyl-L-alanyl-D-glutamate:meso- diaminopimelate ligase | silent codon 91 | HE_50+ |
| *lpd*‡ | b0116 | lipoamide dehydrogenase, E3 component is part ofthree enzyme complexes | silent codon 294 | HE_100+ |
| *-secF*->/****-yajD->*** | b0409/b0410 | SecYEG protein translocase auxillary subunit/conserved protein | T8 -> T9 | HE_50+ |
| ***<-purE-****/<-lpxH*-* | b0523/b0524 | N5-carboxyaminoimidazole ribonucleotide mutase/UDP-2,3-diacylglucosamine pyrophosphatase | A -> G | HE_100+ |
| *-ubiF**‡*->/<-glnX-* | b0662/b0664 | 2-octaprenyl-3-methyl-6-methoxy-1,4-benzoquinol oxygenase/tRNA-Gln(CUG) | A -> G | HE_150 |
| *-nagE->/****-glnS*->*** | b0679/b0680 | fused N-acetyl glucosamine specific PTS enzyme: IIC, IIB, and IIA components/glutamyl-tRNA synthetase | T9 -> T8 | HE_150 |
| *-sdhB->/****-sucA****‡***->*** | b0724/**b0726** | succinate dehydrogenase, FeS subunit/2-oxoglutarate decarboxylase, thiamin-requiring | G -> A | HE_50+ |
| *-mngB->/****-cydA*->*** | b0732/b0733 | alpha-mannosidase/cytochrome d terminal oxidase, subunit I | A -> C | HE_50+ |
| *kdsB* | b0918 | 3-deoxy-manno-octulosonate cytidylyltransferase | silent codon 127 | HE_100+ |
| *-mviM->/****-yceN*->*** | b1068/b1069 | predicted oxidoreductase with NAD(P)-binding Rossmann-fold domain/predicted inner membrane protein | G -> A | HE_50+ |
| *-flgL->/<-rne*-* | b1083/b1084 | flagellar hook-filament junction protein/RNase E | A6 -> A7 | HE_100+ |
| ***<-ydhL****‡***-****/****-ydhM->*** | b1648/b1649 | conserved protein/predicted DNA-binding transcriptional regulator | A -> C | HE_50+ |
| ***<-thrS*-/-arpB->*** | b1719/b4494 | threonyl-tRNA synthetase/putative ankyrin repeat protein B, N-terminal fragment | -C | HE_100+ |
| *-yejH->/****-rplY****‡***->*** | b2184/b2185 | predicted ATP-dependent helicase/50S ribosomal subunit protein L25 | T -> C | HE_50+ |
| *accD* | b2316 | acetyl-CoA carboxylase, beta (carboxyltransferase) subunit | silent codon 298 | HE_50+ |
| *guaA*‡ | b2507 | GMP synthetase (glutamine aminotransferase) | silent codon 378 | HE_50+ |
| *ispG* | b2515 | 1-hydroxy-2-methyl-2-(E)-butenyl 4-diphosphate synthase | silent codon 366 | HE_50+ |
| ***<-rnc*-****/<-lepB*-* | **b2567**/b2568 | RNase III/leader peptidase (signal peptidase I) | C -> T | HE_150 |
| *pyrG* | b2780 | CTP synthetase | silent codon 312 | HE_50+ |
| ***<-ppdA-****/<-thyA**‡*-* | b2826/b2827 | conserved protein/thymidylate synthetase | A7 -> A8 | HE_50+ |
| *-ygeF*->/****-ygeG****‡***->*** | b2850/b2851 | predicted protein/predicted chaperone | A6 -> A5 | HE_50+ |
| *prfB* | **b2891** | peptide chain release factor RF-2 | silent codon 242 | HE_50+ |
| *-ygfZ**‡*->/<-yqfA-* | **b2898**/b2899 | predicted folate-dependent regulatory protein/predicted oxidoreductase, inner membrane subunit | G -> A | HE_50+ |
| ***<-ribB*-/-yqiC->*** | **b3041**/b3042 | 3,4-dihydroxy-2-butanone-4-phosphate synthase/conserved protein | C -> A | HE_50+ |
| *-fadH->/<-ygjM**‡*-* | b3081/b3082 | 2,4-dienoyl-CoA reductase, NADH and FMN-linked/predicted DNA-binding transcriptional regulator | G -> C (50 %) | HE_100^@^ |
| ***<-sspA-****/<-rpsI*-* | b3229/**b3230** | stringent starvation protein A/30S ribosomal subunit protein S9 | A2 -> A3 | HE_150 |
| *def* | b3287 | peptide deformylase | silent codon 103 | HE_50+ |
| *rpsS* | b3316 | 30S ribosomal subunit protein S19 | silent codon 47 | HE_50+ |
| *yrfF* | b3398 | predicted inner membrane protein | silent codon 684 | HE_100+ |
| *-insK->/<-glyS*-* | b3558/**b3559** | IS150 conserved protein InsB/glycine tRNA synthetase, beta subunit | T -> C | HE_50+ |
| *gpsA* | b3608 | glycerol-3-phosphate dehydrogenase (NAD+) | silent codon 201 | HE_50+ |
| *dfp* | b3639 | fused 4'-phosphopantothenoylcysteine decarboxylase; phosphopantothenoylcysteine synthetase, FMN-binding | silent codon 196 | HE_100+ |
| *ubiD*‡ | b3843 | 3-octaprenyl-4-hydroxybenzoate decarboxylase | silent codon 2 | HE_50+ |
| *polA*‡ | **b3863** | DNA polymerase I | silent codon 650 | HE_150 |
| ***<-hslV-****/<-ftsN**‡*-* | b3932/b3933 | peptidase component of the HslUV protease/essential cell division protein | G -> T | HE_50+ |
| *murB* | b3972 | UDP-N-acetylenolpyruvoylglucosamine reductase, FAD-binding | silent codon 217 | HE_50+ |
| ***<-coaA*-/-thrU*->*** | **b3974**/b3976 | pantothenate kinase/tRNA-Thr(UGU) | +T | Parental DH10B+ |
| *-rpoC*->/****-htrC->*** | b3988/b3989 | RNA polymerase, beta prime subunit/heat shock protein | T7 -> T6 | HE_150 |
| *groS* | b4142 | Cpn10 chaperonin GroES, small subunit of GroESL | silent codon 6 | HE_150 |
| *valS* | b4258 | valyl-tRNA synthetase | silent codon 407 | HE_50+ |
| ***<-tdcG-****/<-tdcF**‡*-* | b4471/b3113 | L-serine dehydratase 3/predicted L-PSP (mRNA) endoribonuclease | A -> G | HE_50+ |

‡The essentiality of this gene is unclear, there are conflicting results from essential gene studies.

^@^Mutation reverted after this strain/isolate

**Supplementary Table S6.** Intergenic and silent mutations in or around heat shock genes. For intergenic mutations, the essential genes are indicated with asterisks, arrows surrounding genes indicate their directions and genes in bold have the insertion upstream of their start codons, meaning the insertions may affect the transcriptional or translational regulation of the genes. The b number is shown in bold if the melting temperature of the protein^33^ is below 52 °C (≤5 °C above the maximum LB broth growth temperature of HE_150). All mutations are at >96 % frequency except for *lipB*.

| Gene(s) | B number(s) | Product(s) | Mutation(s) | Strain(s) |
| --- | --- | --- | --- | --- |
| ***<-yaaI-****/****-dnaK*->*** | b0013/b0014 | predicted protein/chaperone Hsp70, co-chaperone with DnaJ | G -> A | HE_50+ |
| *insL-1* | b0016 | IS186/IS421 transposase | silent codon 101 | Parental DH10B ^@^ |
| *hepA* | b0059 | RNA polymerase-associated helicase protein (pseudogene) | silent codon 252 and 528 | HE_50+ |
| *hepA* | b0059 | RNA polymerase-associated helicase protein (pseudogene) | silent codon 110 | HE_100+ |
| ***<-yafC-****/****-yafD*->*** | b0208/b0209 | predicted DNA-binding transcriptional regulator/conserved protein | C -> T | HE_50+ |
| *-cof->/****-ybaO*#->*** | b0446/b0447 | thiamin pyrimidine pyrophosphate hydrolase/predicted DNA-binding transcriptional regulator | C -> T, G -> T | HE_50+ |
| *lipB* | b0630 | lipoyl-protein ligase | silent codon 173 (in one of two copies) | HE_50+ |
| *-nagE->/****-glnS*->*** | b0679/b0680 | fused N-acetyl glucosamine specific PTS enzyme: IIC, IIB, and IIA components/glutamyl-tRNA synthetase | T9 -> T8 | HE_150 |
| ***<-solA-****/<-bssS*-* | b1059/b1060 | N-methyltryptophan oxidase, FAD-binding/regulator of biofilm formation | A -> T | HE_100+ |
| *dgsA* | b1594 | DNA-binding transcriptional repressor | silent codon 396 | HE_50+ |
| ***<-htpX*-****/<-prc-* | b1829/b1830 | predicted endopeptidase/carboxy-terminal protease for penicillin-binding protein 3 | A -> T | HE_100+ |
| *-narP*->/<-ccmH-* | **b2193**/b2194 | DNA-binding response regulator in two-component regulatory system with NarQ or NarX/heme lyase, CcmH subunit | G -> T | HE_50+ |
| *recA* | b2699 | DNA recombination/repair protein RecA | silent codon 331 | HE_50+ |
| ***<-ygaD*-****/<-mltB-* | b2700/b2701 | conserved protein/membrane-bound lytic murein transglycosylase B | C6 -> C5 | HE_50+ |
| ***<-yidE-****/<-ibpB*-* | b3685/b3686 | predicted transporter/heat shock chaperone | C -> T | HE_150 |
| ***<-hslV*-****/<-ftsN-* | b3932/b3933 | peptidase component of the HslUV protease/essential cell division protein | G -> T | HE_50+ |
| *groS* | b4142 | Cpn10 chaperonin GroES, small subunit of GroESL | silent codon 6 | HE_150 |
| *mutL* | b4170 | methyl-directed mismatch repair protein | silent codon 69 | HE_50+ |
| *valS* | b4258 | valyl-tRNA synthetase | silent codon 407 | HE_50+ |

^@^Mutation reverted after this strain/isolate

#Transcription of *ybaO* was significantly elevated in response to σ^32^ overexpression, but unlike with other genes in this table, a σ^32^-dependent promoter was not identified^17^.

**Supplementary Table S7.** Fixed mutations that were lost over the evolution to heat. These mutations were either present in the starting parental DH10B strain (insL-1) or arose at 100 % frequency during the heat evolution process (the remaining mutations). Upon sequencing of later isolates, these mutations were lost and the sequence at these loci became the wild-type DH10B reference sequence.

| Gene(s) | B number | Product(s) | Mutation(s) | Reverted |
| --- | --- | --- | --- | --- |
| *insL-1* | b0016 | IS186/IS421 transposase | silent codon 101 | between parental DH10B and HE_50 |
| *ycbF* | b0944 | predicted periplasmic pilin chaperone | frameshift at codon 72 | between HE_100 and HE_150 |
| *yihA/engB* | b3865 | GTP-binding protein | A30S | between HE_50 and HE_100 |
| *yihO* | b3876 | predicted transporter | D27N | between HE_100 and HE_150 |

**Supplementary Table S8.** Nonfixed mutations that reverted over the evolution to heat. For intergenic mutations, arrows surrounding genes indicate their directions and genes in bold have the insertion upstream of their start codons, meaning the insertions may affect the transcriptional and/or translational regulation of the genes. The b number is shown in bold if the melting temperature of the protein^33^ is below 52 °C (≤5 °C above the maximum LB broth growth temperature of HE_150). The percent frequency of each mutation is shown in brackets in the Mutation(s) column. When this is shown as a range it indicates the range of percent frequencies in the multiple strains the mutation is in.

| Gene(s) | B number(s) | Product(s) | Mutation(s) | Reverted |
| --- | --- | --- | --- | --- |
| ***<-yagI-****/<-argF-* | b0272/b0273 | CP4-6 prophage; DNA-binding transcriptional repressor/CP4-6 prophage; ornithine carbamoyltransferase 2, chain F | G -> C (31 %) | between parental DH10B and HE_50 |
| *ychS* | b1228 | predicted protein | K29E (31 %) | between HE_100 and HE_150 |
| *-dmsD->/****- clcB->*** | b1591/b1592 | twin-argninine leader-binding protein for DmsA and TorA/  predicted voltage-gated chloride channel | C -> T (40 %) | between parental DH10B and HE_50 |
| *yeeO* | b1985 | predicted multidrug efflux system | truncation at codon 2 (27 %) | between parental DH10B and HE_50 |
| *-fadH->/<-ygjM-* | b3081/b3082 | 2,4-dienoyl-CoA reductase, NADH and FMN-linked/predicted DNA-binding transcriptional regulator | G -> C (50 %) | between HE_100 and HE_150 |
| *tdcD* | b3115 | propionate kinase/acetate kinase C, anaerobic | silent codon 60 (28 %) | between HE_100 and HE_150 |
| *obgE* | **b3183** | GTPase involved in cell partitioning and DNA repair | S75T (39 %) | between HE_100 and HE_150 |
| ***<-yhfG-****/<-ppiA-* | b3362/b3363 | predicted protein/peptidyl-prolyl cis-trans isomerase A (rotamase A) | C -> G (81 %) | between HE_100 and HE_150 |
| ***<-yhfG-****/<-ppiA-* | b3362/b3363 | predicted protein/peptidyl-prolyl cis-trans isomerase A (rotamase A) | CA -> GG (≥50 %) | between HE_100 and HE_150 |

**Supplementary Table S9.** Changes in the predicted translation initiation rate (TIR; https://www.denovodna.com/software/predict_rbs_calculator)^20^ of genes with mutations near the start codon. Genes with known transcript start sites and mutations near the start codon were analyzed. For *ubiD*, two different transcripts were used and the values in brackets correspond to the second transcript. The TIR after mutation is shown in bold if the mutation changed the TIR.

| Gene | B number | 5' UTR length | TIR in DH10B (au) | Strain(s) with mutation | TIR after mutation (au) |
| --- | --- | --- | --- | --- | --- |
| *yafD* | b0209 | 58 | 316.7 | HE_50+ | **416.75** |
| *sucA* | b0726 | 120 | 252.35 | HE_50+ | **552.2** |
| *fabH* | b1091 | 243 | 252.11 | HE_150 | **79.66** |
| *rssB* | b1235 | 22 | 1015.04 | HE_50+ | 1015.04 |
| *ribB* | b3041 | 252 | 807.6 | HE_50+ | **594.68** |
| *selA* | b3591 | 48 | 2134.48 | HE_50+ | 2134.48 |
| *ubiD* | b3843 | 18 (179) | 195.47 (167.21) | HE_50+ | **316.39 (199.29)** |
| *glpF* | b3927 | 71 | 8201.32 | HE_100+ | 8201.32 |
| *lexA* | b4043 | 28 | 2323.04 | HE_50+ | 2323.04 |
| *groS* | b4142 | 72 | 17753.13 | HE_150 | **24547.11** |

**Supplementary Table S10.** Gene Ontology biological processes overrepresented and underrepresented in the protein expression categories. There were no overrepresentations nor underrepresentations for the restored protein expression category.

| Gene  expression  category | GO biological process  complete | Escherichia  coli  REFLIST  (4392) | upload | upload  (expected) | upload  (over/under) | upload  (fold  Enrichment) | upload  (raw  P-value) | upload  (FDR) |
| --- | --- | --- | --- | --- | --- | --- | --- | --- |
| Unrestored | protein folding  (GO:0006457) | 57 | 6 | 0.42 | + | 14.45 | 4.44E-06 | 1.37E-02 |
| Reinforced | Mo-molybdopterin  cofactor biosynthetic  process (GO:0006777) | 14 | 3 | 0.03 | + | 94.11 | 5.65E-06 | 8.68E-03 |
| Novel | glycolytic process  (GO:0006096) | 19 | 5 | 0.43 | + | 11.67 | 1.43E-04 | 2.20E-02 |
| Novel | carboxylic acid catabolic  process (GO:0046395) | 203 | 14 | 4.58 | + | 3.06 | 2.43E-04 | 3.56E-02 |
| Novel | amino acid metabolic  process (GO:0006520) | 261 | 17 | 5.88 | + | 2.89 | 9.54E-05 | 1.83E-02 |
| Novel | phosphate-containing  compound metabolic  process (GO:0006796) | 430 | 23 | 9.69 | + | 2.37 | 1.14E-04 | 2.05E-02 |
| Novel | regulation of cellular  process (GO:0050794) | 592 | 2 | 13.34 | - | 0.15 | 1.39E-04 | 2.25E-02 |

**Supplementary Table S11.** Lists of proteins in each protein expression category from overrepresented and underrepresented Gene Ontology biological processes.

| Unrestored | | | |
| --- | --- | --- | --- |
|  | protein folding (GO:0006457) | | |
|  |  | ivy | Inhibitor of vertebrate lysozyme;ivy;;orthologs |
|  |  | degP | Periplasmic serine endoprotease DegP;degP;PTN001117973;orthologs |
|  |  | dnaK | Chaperone protein DnaK;dnaK;PTN000452647;orthologs |
|  |  | dppA | Dipeptide-binding protein;dppA;PTN000767719;orthologs |
|  |  | clpB | Chaperone protein ClpB;clpB;PTN000181415;orthologs |
|  |  | htpG | Chaperone protein HtpG;htpG;PTN000163845;orthologs |
| Reinforced | | | |
|  | Mo-molybdopterin cofactor biosynthetic process (GO:0006777) | | |
|  |  | moaE | Molybdopterin synthase catalytic subunit;moaE;PTN000600974;orthologs |
|  |  | moaC | Cyclic pyranopterin monophosphate synthase;moaC;;orthologs |
|  |  | moaB | Molybdenum cofactor biosynthesis protein B;moaB;PTN000022877;orthologs |
| Novel | | | |
|  | glycolytic process (GO:0006096) | | |
|  |  | pfkB | ATP-dependent 6-phosphofructokinase isozyme 2;pfkB;PTN000833101;orthologs |
|  |  | fbaB | Fructose-bisphosphate aldolase class 1;fbaB;PTN005360439;orthologs |
|  |  | pykF | Pyruvate kinase I;pykF;PTN000212867;orthologs |
|  |  | tpiA | Triosephosphate isomerase;tpiA;PTN001079909;orthologs |
|  |  | glk | Glucokinase;glk;PTN001253784;orthologs |
|  | carboxylic acid catabolic process (GO:0046395) | | |
|  |  | yagE | Putative 2-dehydro-3-deoxy-D-gluconate aldolase YagE;yagE;PTN000252865;orthologs |
|  |  | nagB | Glucosamine-6-phosphate deaminase;nagB;PTN001345265;orthologs |
|  |  | putA | Bifunctional protein PutA;putA;PTN000192505;orthologs |
|  |  | tnaA | Tryptophanase;tnaA;PTN000794797;orthologs |
|  |  | garR | 2-hydroxy-3-oxopropionate reductase;garR;PTN000541514;orthologs |
|  |  | fadA | 3-ketoacyl-CoA thiolase FadA;fadA;PTN000432476;orthologs |
|  |  | dadX | Alanine racemase, catabolic;dadX;PTN000775518;orthologs |
|  |  | aldA | Lactaldehyde dehydrogenase;aldA;PTN001364926;orthologs |
|  |  | nanK | N-acetylmannosamine kinase;nanK;PTN001056113;orthologs |
|  |  | gabD | Succinate-semialdehyde dehydrogenase [NADP(+)] GabD;gabD;PTN001364927;orthologs |
|  |  | tusA | Sulfur carrier protein TusA;tusA;PTN002116241;orthologs |
|  |  | gabT | 4-aminobutyrate aminotransferase GabT;gabT;PTN000944969;orthologs |
|  |  | poxB | Pyruvate dehydrogenase [ubiquinone];poxB;PTN000438957;orthologs |
|  |  | dadA | D-amino acid dehydrogenase;dadA;PTN001014654;orthologs |
|  | amino acid metabolic process (GO:0006520) | | |
|  |  | argG | Argininosuccinate synthase;argG;PTN000903732;orthologs |
|  |  | metG | Methionine--tRNA ligase;metG;PTN000942565;orthologs |
|  |  | putA | Bifunctional protein PutA;putA;PTN000192505;orthologs |
|  |  | tnaA | Tryptophanase;tnaA;PTN000794797;orthologs |
|  |  | dadX | Alanine racemase, catabolic;dadX;PTN000775518;orthologs |
|  |  | thrC | Threonine synthase;thrC;PTN000034359;orthologs |
|  |  | serA | D-3-phosphoglycerate dehydrogenase;serA;PTN000108067;orthologs |
|  |  | aldA | Lactaldehyde dehydrogenase;aldA;PTN001364926;orthologs |
|  |  | nadE | NH(3)-dependent NAD(+) synthetase;nadE;;orthologs |
|  |  | gabD | Succinate-semialdehyde dehydrogenase [NADP(+)] GabD;gabD;PTN001364927;orthologs |
|  |  | guaA | GMP synthase [glutamine-hydrolyzing];guaA;PTN000938422;orthologs |
|  |  | tusA | Sulfur carrier protein TusA;tusA;PTN002116241;orthologs |
|  |  | gabT | 4-aminobutyrate aminotransferase GabT;gabT;PTN000944969;orthologs |
|  |  | folA | Dihydrofolate reductase;folA;PTN001356377;orthologs |
|  |  | hisG | ATP phosphoribosyltransferase;hisG;PTN001468430;orthologs |
|  |  | metH | Methionine synthase;metH;PTN000473136;orthologs |
|  |  | dadA | D-amino acid dehydrogenase;dadA;PTN001014654;orthologs |
|  | phosphate-containing compound metabolic process (GO:0006796) | | |
|  |  | pgl | 6-phosphogluconolactonase;pgl;PTN001251288;orthologs |
|  |  | pfkB | ATP-dependent 6-phosphofructokinase isozyme 2;pfkB;PTN000833101;orthologs |
|  |  | agp | Glucose-1-phosphatase;agp;PTN001696355;orthologs |
|  |  | glgP | Glycogen phosphorylase;glgP;PTN000895158;orthologs |
|  |  | glpK | Glycerol kinase;glpK;PTN000810551;orthologs |
|  |  | pncC | Nicotinamide-nucleotide amidohydrolase PncC;pncC;PTN000358658;orthologs |
|  |  | speG | Spermidine N(1)-acetyltransferase;speG;PTN000327284;orthologs |
|  |  | dhaK | PEP-dependent dihydroxyacetone kinase, dihydroxyacetone-binding subunit DhaK;dhaK;PTN002008120;orthologs |
|  |  | ushA | Protein UshA;ushA;PTN000171653;orthologs |
|  |  | nudE | ADP compounds hydrolase NudE;nudE;PTN000217390;orthologs |
|  |  | pykF | Pyruvate kinase I;pykF;PTN000212867;orthologs |
|  |  | tpiA | Triosephosphate isomerase;tpiA;PTN001079909;orthologs |
|  |  | mak | Fructokinase;mak;PTN001056115;orthologs |
|  |  | grxC | Glutaredoxin 3;grxC;PTN000018845;orthologs |
|  |  | glk | Glucokinase;glk;PTN001253784;orthologs |
|  |  | nadE | NH(3)-dependent NAD(+) synthetase;nadE;;orthologs |
|  |  | nanK | N-acetylmannosamine kinase;nanK;PTN001056113;orthologs |
|  |  | gpsA | Glycerol-3-phosphate dehydrogenase [NAD(P)+];gpsA;PTN001366299;orthologs |
|  |  | tdk | Thymidine kinase;tdk;PTN000893565;orthologs |
|  |  | guaA | GMP synthase [glutamine-hydrolyzing];guaA;PTN000938422;orthologs |
|  |  | tusA | Sulfur carrier protein TusA;tusA;PTN002116241;orthologs |
|  |  | manX | PTS system mannose-specific EIIAB component;manX;PTN002138935;orthologs |
|  |  | rbsK | Ribokinase;rbsK;PTN000062092;orthologs |
|  | regulation of cellular process (GO:0050794) | | |
|  |  | putA | Bifunctional protein PutA;putA;PTN000192505;orthologs |
|  |  | dps | DNA protection during starvation protein;dps;PTN002016965;orthologs |

**Supplementary Table S12.** Gene Ontology biological processes overrepresented and underrepresented for proteins at significantly different abundances in HE_150 at 42.5°C compared to DH10B at 37 °C. All values were calculated by the Gene Ontology Database using a Fisher’s Exact Test with a Calculate False Discovery Rate Correction.

| Lower or higher in HE_150 at 42.5°C compared to DH10B at 37 °C | GO biological process  complete | Escherichia  coli  REFLIST  (4392) | upload | upload  (expected) | upload  (over/under) | upload  (fold  Enrichment) | upload  (raw  P-value) | upload  (FDR) |
| --- | --- | --- | --- | --- | --- | --- | --- | --- |
| Lower | aspartate metabolic process (GO:0006531) | 8 | 4 | 0.24 | + | 16.64 | 2.86E-04 | 3.38E-02 |
| Lower | pyruvate metabolic process (GO:0006090) | 38 | 7 | 1.14 | + | 6.13 | 2.76E-04 | 3.39E-02 |
| Lower | protein homotetramerization (GO:0051289) | 48 | 8 | 1.44 | + | 5.55 | 1.85E-04 | 2.59E-02 |
| Lower | carbohydrate catabolic process (GO:0016052) | 138 | 17 | 4.15 | + | 4.1 | 1.85E-06 | 7.09E-04 |
| Lower | amino acid catabolic process (GO:0009063) | 82 | 10 | 2.46 | + | 4.06 | 2.99E-04 | 3.41E-02 |
| Lower | monosaccharide metabolic process (GO:0005996) | 109 | 12 | 3.28 | + | 3.66 | 1.79E-04 | 2.62E-02 |
| Lower | alpha-amino acid biosynthetic process (GO:1901607) | 120 | 13 | 3.61 | + | 3.6 | 1.11E-04 | 2.00E-02 |
| Lower | response to oxidative stress (GO:0006979) | 102 | 11 | 3.07 | + | 3.59 | 3.95E-04 | 4.34E-02 |
| Lower | regulation of cellular process (GO:0050794) | 592 | 4 | 17.79 | - | 0.22 | 1.13E-04 | 1.82E-02 |
| Lower | nucleic acid metabolic process (GO:0090304) | 515 | 2 | 15.48 | - | 0.13 | 3.54E-05 | 9.06E-03 |
| Higher | regulation of protein catabolic process (GO:0042176) | 5 | 3 | 0.1 | + | 31 | 3.45E-04 | 2.87E-02 |
| Higher | histidine biosynthetic process (GO:0000105) | 10 | 4 | 0.19 | + | 20.67 | 1.05E-04 | 1.19E-02 |
| Higher | Mo-molybdopterin cofactor biosynthetic process (GO:0006777) | 14 | 5 | 0.27 | + | 18.45 | 2.07E-05 | 3.97E-03 |
| Higher | protein folding (GO:0006457) | 57 | 7 | 1.1 | + | 6.35 | 1.79E-04 | 1.72E-02 |
| Higher | response to heat (GO:0009408) | 68 | 8 | 1.32 | + | 6.08 | 7.99E-05 | 9.82E-03 |
| Higher | transmembrane transport (GO:0055085) | 645 | 1 | 12.48 | - | 0.08 | 5.67E-05 | 7.25E-03 |

**Supplementary Table S13.** Lists of proteins at significantly different abundances in HE_150 at 42.5°C compared to DH10B at 37 °C from overrepresented and underrepresented Gene Ontology biological processes.

| At significantly lower abundance in HE_150 at 42.5 °C than DH10B at 37 °C | | | |
| --- | --- | --- | --- |
|  | aspartate metabolic process (GO:0006531) | | |
|  |  | tyrB | Aromatic-amino-acid aminotransferase;tyrB;PTN000222934;orthologs |
|  |  | aspA | Aspartate ammonia-lyase;aspA;PTN000893832;orthologs |
|  |  | nadE | NH(3)-dependent NAD(+) synthetase;nadE;;orthologs |
|  |  | ilvE | Branched-chain-amino-acid aminotransferase;ilvE;PTN001371641;orthologs |
|  | pyruvate metabolic process (GO:0006090) | | |
|  |  | pfkB | ATP-dependent 6-phosphofructokinase isozyme 2;pfkB;PTN000833101;orthologs |
|  |  | dhaK | PEP-dependent dihydroxyacetone kinase, dihydroxyacetone-binding subunit DhaK;dhaK;PTN002008120;orthologs |
|  |  | fbaB | Fructose-bisphosphate aldolase class 1;fbaB;PTN005360439;orthologs |
|  |  | pykF | Pyruvate kinase I;pykF;PTN000212867;orthologs |
|  |  | tpiA | Triosephosphate isomerase;tpiA;PTN001079909;orthologs |
|  |  | glk | Glucokinase;glk;PTN001253784;orthologs |
|  |  | poxB | Pyruvate dehydrogenase [ubiquinone];poxB;PTN000438957;orthologs |
|  | protein homotetramerization (GO:0051289) | | |
|  |  | ppnN | Pyrimidine/purine nucleotide 5'-monophosphate nucleosidase;ppnN;PTN001581542;orthologs |
|  |  | glgC | Glucose-1-phosphate adenylyltransferase;glgC;PTN001475582;orthologs |
|  |  | aspA | Aspartate ammonia-lyase;aspA;PTN000893832;orthologs |
|  |  | rfbA | Glucose-1-phosphate thymidylyltransferase 1;rfbA;PTN001096497;orthologs |
|  |  | gabD | Succinate-semialdehyde dehydrogenase [NADP(+)] GabD;gabD;PTN001364927;orthologs |
|  |  | ppc | Phosphoenolpyruvate carboxylase;ppc;PTN001253963;orthologs |
|  |  | srlD | Sorbitol-6-phosphate 2-dehydrogenase;srlD;PTN001292941;orthologs |
|  |  | speA | Biosynthetic arginine decarboxylase;speA;PTN000896467;orthologs |
|  | carbohydrate catabolic process (GO:0016052) | | |
|  |  | pfkB | ATP-dependent 6-phosphofructokinase isozyme 2;pfkB;PTN000833101;orthologs |
|  |  | agp | Glucose-1-phosphatase;agp;PTN001696355;orthologs |
|  |  | glgP | Glycogen phosphorylase;glgP;PTN000895158;orthologs |
|  |  | glpK | Glycerol kinase;glpK;PTN000810551;orthologs |
|  |  | yagH | Putative beta-xylosidase;yagH;PTN000531949;orthologs |
|  |  | garR | 2-hydroxy-3-oxopropionate reductase;garR;PTN000541514;orthologs |
|  |  | dhaK | PEP-dependent dihydroxyacetone kinase, dihydroxyacetone-binding subunit DhaK;dhaK;PTN002008120;orthologs |
|  |  | fbaB | Fructose-bisphosphate aldolase class 1;fbaB;PTN005360439;orthologs |
|  |  | pykF | Pyruvate kinase I;pykF;PTN000212867;orthologs |
|  |  | aldA | Lactaldehyde dehydrogenase;aldA;PTN001364926;orthologs |
|  |  | tpiA | Triosephosphate isomerase;tpiA;PTN001079909;orthologs |
|  |  | glk | Glucokinase;glk;PTN001253784;orthologs |
|  |  | galE | UDP-glucose 4-epimerase;galE;PTN000041930;orthologs |
|  |  | uxaC | Uronate isomerase;uxaC;PTN000763890;orthologs |
|  |  | srlD | Sorbitol-6-phosphate 2-dehydrogenase;srlD;PTN001292941;orthologs |
|  |  | rbsK | Ribokinase;rbsK;PTN000062092;orthologs |
|  |  | fucO | Lactaldehyde reductase;fucO;PTN002608726;orthologs |
|  | amino acid catabolic process (GO:0009063) | | |
|  |  | putA | Bifunctional protein PutA;putA;PTN000192505;orthologs |
|  |  | tnaA | Tryptophanase;tnaA;PTN000794797;orthologs |
|  |  | aspA | Aspartate ammonia-lyase;aspA;PTN000893832;orthologs |
|  |  | dadX | Alanine racemase, catabolic;dadX;PTN000775518;orthologs |
|  |  | aldA | Lactaldehyde dehydrogenase;aldA;PTN001364926;orthologs |
|  |  | gabD | Succinate-semialdehyde dehydrogenase [NADP(+)] GabD;gabD;PTN001364927;orthologs |
|  |  | gabT | 4-aminobutyrate aminotransferase GabT;gabT;PTN000944969;orthologs |
|  |  | gadB | Glutamate decarboxylase beta;gadB;PTN000945882;orthologs |
|  |  | speA | Biosynthetic arginine decarboxylase;speA;PTN000896467;orthologs |
|  |  | dadA | D-amino acid dehydrogenase;dadA;PTN001014654;orthologs |
|  | monosaccharide metabolic process (GO:0005996) | | |
|  |  | pgl | 6-phosphogluconolactonase;pgl;PTN001251288;orthologs |
|  |  | agp | Glucose-1-phosphatase;agp;PTN001696355;orthologs |
|  |  | dhaK | PEP-dependent dihydroxyacetone kinase, dihydroxyacetone-binding subunit DhaK;dhaK;PTN002008120;orthologs |
|  |  | aldA | Lactaldehyde dehydrogenase;aldA;PTN001364926;orthologs |
|  |  | tpiA | Triosephosphate isomerase;tpiA;PTN001079909;orthologs |
|  |  | dkgB | 2,5-diketo-D-gluconic acid reductase B;dkgB;PTN000920826;orthologs |
|  |  | ppc | Phosphoenolpyruvate carboxylase;ppc;PTN001253963;orthologs |
|  |  | galE | UDP-glucose 4-epimerase;galE;PTN000041930;orthologs |
|  |  | uxaC | Uronate isomerase;uxaC;PTN000763890;orthologs |
|  |  | rbsK | Ribokinase;rbsK;PTN000062092;orthologs |
|  |  | fucO | Lactaldehyde reductase;fucO;PTN002608726;orthologs |
|  |  | xylF | D-xylose-binding periplasmic protein;xylF;PTN000769033;orthologs |
|  | alpha-amino acid biosynthetic process (GO:1901607) | | |
|  |  | argG | Argininosuccinate synthase;argG;PTN000903732;orthologs |
|  |  | tyrB | Aromatic-amino-acid aminotransferase;tyrB;PTN000222934;orthologs |
|  |  | carA | Carbamoyl-phosphate synthase small chain;carA;PTN000892052;orthologs |
|  |  | putA | Bifunctional protein PutA;putA;PTN000192505;orthologs |
|  |  | carB | Carbamoyl-phosphate synthase large chain;carB;PTN000150349;orthologs |
|  |  | dadX | Alanine racemase, catabolic;dadX;PTN000775518;orthologs |
|  |  | ilvD | Dihydroxy-acid dehydratase;ilvD;PTN001464129;orthologs |
|  |  | thrC | Threonine synthase;thrC;PTN000034359;orthologs |
|  |  | gabT | 4-aminobutyrate aminotransferase GabT;gabT;PTN000944969;orthologs |
|  |  | folA | Dihydrofolate reductase;folA;PTN001356377;orthologs |
|  |  | ilvE | Branched-chain-amino-acid aminotransferase;ilvE;PTN001371641;orthologs |
|  |  | metH | Methionine synthase;metH;PTN000473136;orthologs |
|  |  | asnA | Aspartate--ammonia ligase;asnA;PTN001247838;orthologs |
|  | response to oxidative stress (GO:0006979) | | |
|  |  | osmC | Peroxiredoxin OsmC;osmC;PTN002138792;orthologs |
|  |  | ygiW | Protein YgiW;ygiW;PTN002195215;orthologs |
|  |  | acnA | Aconitate hydratase A;acnA;PTN000912601;orthologs |
|  |  | putA | Bifunctional protein PutA;putA;PTN000192505;orthologs |
|  |  | grxC | Glutaredoxin 3;grxC;PTN000018845;orthologs |
|  |  | katE | Catalase HPII;katE;PTN000157377;orthologs |
|  |  | btuE | Thioredoxin/glutathione peroxidase BtuE;btuE;PTN000904507;orthologs |
|  |  | fumC | Fumarate hydratase class II;fumC;PTN000154580;orthologs |
|  |  | yajQ | UPF0234 protein YajQ;yajQ;PTN000767169;orthologs |
|  |  | wrbA | NAD(P)H dehydrogenase (quinone);wrbA;PTN001254453;orthologs |
|  |  | katG | Catalase-peroxidase;katG;PTN001254519;orthologs |
|  | regulation of cellular process (GO:0050794) | | |
|  |  | putA | Bifunctional protein PutA;putA;PTN000192505;orthologs |
|  |  | thyA | Thymidylate synthase;thyA;PTN000900110;orthologs |
|  |  | dps | DNA protection during starvation protein;dps;PTN002016965;orthologs |
|  |  | cspD | Cold shock-like protein CspD;cspD;PTN000166685;orthologs |
|  | nucleic acid metabolic process (GO:0090304) | | |
|  |  | yegP | UPF0339 protein YegP;yegP;PTN002221064;orthologs |
|  |  | ligA | DNA ligase;ligA;PTN000121749;orthologs |
| At significantly higher abundance in HE_150 at 42.5 °C than DH10B at 37 °C | | | |
|  | regulation of protein catabolic process (GO:0042176) | | |
|  |  | iraP | Anti-adapter protein IraP;iraP;;orthologs |
|  |  | sspB | Stringent starvation protein B;sspB;PTN002206208;orthologs |
|  |  | hslU | ATP-dependent protease ATPase subunit HslU;hslU;PTN000137399;orthologs |
|  | histidine biosynthetic process (GO:0000105) | | |
|  |  | hisC | Histidinol-phosphate aminotransferase;hisC;PTN002872500;orthologs |
|  |  | hisF | Imidazole glycerol phosphate synthase subunit HisF;hisF;PTN001082185;orthologs |
|  |  | hisH | Imidazole glycerol phosphate synthase subunit HisH;hisH;PTN001082228;orthologs |
|  |  | hisG | ATP phosphoribosyltransferase;hisG;PTN001468430;orthologs |
|  | Mo-molybdopterin cofactor biosynthetic process (GO:0006777) | | |
|  |  | moaE | Molybdopterin synthase catalytic subunit;moaE;PTN000600974;orthologs |
|  |  | moaC | Cyclic pyranopterin monophosphate synthase;moaC;;orthologs |
|  |  | moaB | Molybdenum cofactor biosynthesis protein B;moaB;PTN000022877;orthologs |
|  |  | tusA | Sulfur carrier protein TusA;tusA;PTN002116241;orthologs |
|  |  | moaD | Molybdopterin synthase sulfur carrier subunit;moaD;PTN002120445;orthologs |
|  | protein folding (GO:0006457) | | |
|  |  | hscB | Co-chaperone protein HscB;hscB;PTN000361340;orthologs |
|  |  | groS | 10 kDa chaperonin;groS;PTN000080810;orthologs |
|  |  | hslO | 33 kDa chaperonin;hslO;PTN000764703;orthologs |
|  |  | clpB | Chaperone protein ClpB;clpB;PTN000181415;orthologs |
|  |  | htpG | Chaperone protein HtpG;htpG;PTN000163845;orthologs |
|  |  | groL | 60 kDa chaperonin;groL;PTN000143676;orthologs |
|  |  | oppA | Periplasmic oligopeptide-binding protein;oppA;PTN000767696;orthologs |
|  | response to heat (GO:0009408) | | |
|  |  | groS | 10 kDa chaperonin;groS;PTN000080810;orthologs |
|  |  | hslO | 33 kDa chaperonin;hslO;PTN000764703;orthologs |
|  |  | hslU | ATP-dependent protease ATPase subunit HslU;hslU;PTN000137399;orthologs |
|  |  | hslV | ATP-dependent protease subunit HslV;hslV;PTN000175847;orthologs |
|  |  | clpB | Chaperone protein ClpB;clpB;PTN000181415;orthologs |
|  |  | htpG | Chaperone protein HtpG;htpG;PTN000163845;orthologs |
|  |  | groL | 60 kDa chaperonin;groL;PTN000143676;orthologs |
|  |  | oppA | Periplasmic oligopeptide-binding protein;oppA;PTN000767696;orthologs |
|  | transmembrane transport (GO:0055085) | | |
|  |  | oppA | Periplasmic oligopeptide-binding protein;oppA;PTN000767696;orthologs |

**Supplementary Methods**

**Proteomics methods as provided by The University of Victoria Genome BC Proteomics Centre**

**(**[**https://www.proteincentre.com/**](https://www.proteincentre.com/)**). Copied with permission.**

**Protein Determination and In-Solution Digestion**

Protein concentration was determined using the bicinchoninic acid (BCA) assay to be: 1. 0.66 μg/μL, 2. 0.57 μg/μL, 3. 0.85 μg/μL, 4. 0.61 μg/μL. 20 μg protein from each sample was precipitated with 10 volumes of acetone overnight at -20°C. Samples were centrifuged at 6,000 x g for 10 min at 10°C. The supernatant was removed and the pellet was resuspended in 40 μL 9 M urea/ 300 mM Tris pH 8.0. The protein samples were reduced by adding dithiothreitol (DTT) in 9 M urea/ 300 mM Tris pH 8.0 to a final concentration of 20 mM DTT, and samples were incubated at 37°C for 30 minutes. To alkylate the samples, iodoacetamide (IAA) in MS-grade water was added to a final concentration of 40 mM IAA, and samples were incubated in the dark at room temperature for 30 minutes. The samples were diluted with 400 μL 100 mM Tris pH 8.0 to reduce the urea concentration to 0.78 M. Modified sequencing grade porcine trypsin (Promega) was reconstituted in 100 mM Tris pH 8.0, trypsin was added to 10:1 substrate: enzyme ratio, and the samples were incubated at 37°C for 18 hours. The samples were acidified with formic acid (FA) to a final concentration of 5% FA, and desalted/concentrated using 200 μL C18 material Stage Tips (Thermo Scientific, catalogue number SP301) following the manufacturers protocol. The peptides were eluted in 40 μL 80% acetonitrile/ 5% FA and speed vacuum concentrated prior to LC- MS/MS analysis.

**LC-MS/MS analysis: Orbitrap Fusion**

The peptide digests ~0.75μg (3μL injection) were separated by on-line reverse phase chromatography using a Thermo Scientific EASY-nLC 1000 system with an Acclaim PepMap100 C18 (100μm I.D., 2 cm length, 5μm, 100Å) reversed-phase pre- column, and an AcclaimPepMap100 C-18 (75μm I.D., 15 cm length, 3μm, 100Å, Thermo Fisher Scientific, San Jose, CA) reversed phase nano-analytical column at a flow rate of 300nl/min. The chromatography system was coupled on-line with an Orbitrap Fusion Tribrid mass spectrometer (Thermo Fisher Scientific, San Jose, CA) equipped with a Nanospray Flex NG source (Thermo Fisher Scientific). Solvents were A: 2% Acetonitrile, 0.1% Formic acid; B: 90% Acetonitrile, 0.1% Formic acid. After a 348 bar (~ 4μL) pre- column equilibration and 348 bar (~ 4μL) nanocolumn equilibration, samples were separated by a 140-minute gradient (0 min: 5%B; 100 min: 25%B; 20 min: 40%B; 10min: 90%B; hold 5 min 90%B); 1min: 100%B; hold 4 min 100%B). The Orbitrap Fusion instrument parameters (Fusion Tune 3.5 software) were as follows for orbitrap (OT-MS) iontrap (IT- MS/MS) with HCD fragmentation: Nano- electrospray ion source with spray voltage 2.65kV, capillary temperature 275 °C. The acquired survey MS1 scan range was 350-1800 m/z profile mode, resolution 120,000 FWHM@200m/z one microscan with Automatic inject time with the Siloxane mass 445.12003 used as lock mass for internal calibration.

Data-dependent acquisition Orbitrap survey spectra were scheduled at least every 3 seconds, with the software determining “Automatic” number of MS/MS acquisitions during this period. The automatic gain control (AGC) target value for FTMS was set to Standard (400,000 counts) and automatic maximum fill time to ensure optimal sensitivity and cycle time. The most intense ions charge state 2-5 exceeding 20,000 counts were selected for HCD MSMS fragmentation in the ion routing multipole. Monoisotopic Precursor Selection (MIPS) was enabled and Dynamic exclusion settings were: repeat count: 2; repeat duration: 5 seconds; exclusion duration: 5 seconds with a 10ppm mass window. The data dependent (ddMS2) IT HCD scan used a quadrupole isolation window of 1.6 Da; Iontrap rapid scan rate, automode normal m/z range, centroid detection, 1 microscan, Auto maximum injection time, AGC target (Standard) 10,000 counts and stepped HCD collision energy of 27,30,33%.

**Data Analysis Parameters**

Protein Sequence Databases

Protein sequence databases for each strain (DH10B, https://www.ncbi.nlm.nih.gov/nuccore/CP110018.1/ and HE150,

(https://www.ncbi.nlm.nih.gov/nuccore/CP110015.1/) were downloaded in FASTA format and combined using Galaxy

v1.2.0 (https://usegalaxy.org/). The two files were merged to produce a single FASTA using the ‘FASTA Merge Files and

Filter Unique Sequences’ tool, where DH10B was used as the primary (reference) strain. Common contaminants were

also added to the final FASTA file used for protein identification.

Protein Identification and Quantification

Proteins in each sample were identified and relative quantitation performed using the software tools FragPipe* (v20.0),

MSFragger (v3.8), and IonQuant (v1.9.8). Search settings included trypsin as the cleavage enzyme with a maximum of

two missed cleavages, N-terminal acetylation and oxidized methionine as variable modifications, and

carbamidomethylation of cysteine as a fixed modification. The output of the database search was filtered to an FDR of

0.01, and default IonQuant settings were used including MaxLFQ and Match between runs.

*References:

i. Kong, A. T., Leprevost, F. V., Avtonomov, D. M., Mellacheruvu, D., & Nesvizhskii, A. I. (2017). MSFragger:ultrafast and comprehensive peptide identification in mass spectrometry–based proteomics. Nature Methods, 14(5), 513-520.
ii. Yu, F., Haynes, S. E., & Nesvizhskii, A. I. (2021). IonQuant enables accurate and sensitive label-free quantification with FDR-controlled match-between-runs. Molecular & Cellular Proteomics, 20.
iii. Teo, G. C., Polasky, D. A., Yu, F., Nesvizhskii, A. I. (2020). A fast deisotoping algorithm and its implementation in the MSFragger search engine. Journal of Proteome Research.
